# Neural mechanisms underlying uninstructed orofacial movements during reward-based learning behaviors

**DOI:** 10.1101/2022.10.28.514159

**Authors:** Wanru Li, Takashi Nakano, Kohta Mizutani, Takanori Matsubara, Masahiro Kawatani, Yasutaka Mukai, Teruko Danjo, Hikaru Ito, Hidenori Aizawa, Akihiro Yamanaka, Carl C. H. Petersen, Junichiro Yoshimoto, Takayuki Yamashita

**Affiliations:** Department of Physiology, Fujita Health University School of Medicine, Toyoake, Japan; Department of Neuroscience II, Research Institute of Environmental Medicine, Nagoya University, Nagoya, Japan; Department of Functional Anatomy & Neuroscience, Graduate School of Medicine, Nagoya University, Nagoya, Japan; Department of Computational Biology, Fujita Health University School of Medicine, Toyoake, Japan; Nara Institute of Science and Technology, Ikoma, Japan; International Center for Brain Science (ICBS), Fujita Health University, Toyoake, Japan; Laboratory for Advanced Brain Functions, Institute for Protein Research, Osaka University, Osaka, Japan; Department of Neurobiology, Graduate School of Biomedical and Health Sciences, Hiroshima University, Hiroshima, Japan; Research Facility Center for Science and Technology, Kagawa University, Kagawa, Japan; Laboratory of Sensory Processing, Brain Mind Institute, Faculty of Life Sciences, École Polytechnique Fédérale de Lausanne (EPFL), Lausanne, Switzerland; Department of Biomedical Data Science, Fujita Health University School of Medicine, Toyoake, Japan

## Abstract

During reward-based learning tasks, animals make orofacial movements that globally influence brain activity at the timings of reward expectation and acquisition. These orofacial movements are not explicitly instructed and typically appear along with goal-directed behaviors. Here we show that reinforcing optogenetic stimulation of midbrain dopamine neurons (oDAS) in mice is sufficient to induce orofacial movements in the whiskers and nose without accompanying goal-directed behaviors. Pavlovian conditioning with a sensory cue and oDAS elicited cue-locked and oDAS aligned orofacial movements, which were distinguishable by a machine learning model. Inhibition or knock-out of dopamine D1 receptors in the nucleus accumbens inhibited oDAS-induced motion but spared cue-locked motion, suggesting differential neural regulation of these two types of orofacial motions. In contrast, inactivation of the whisker primary motor cortex (wM1) abolished both types of orofacial movements. We found specific neuronal populations in wM1 representing either oDAS-aligned or cue-locked whisker movements. Notably, optogenetic stimulation of wM1 neurons successfully replicated these two types of movements. Our results thus suggest that accumbal D1 receptor-dependent and -independent neuronal signals converge in the wM1 for facilitating uninstructed orofacial movements during a reward-based learning task.

## INTRODUCTION

It has long been known that the movements of animals, such as locomotion and whisking, profoundly influence neuronal activity within sensory cortices^1–3^. Recent large-scale neural recordings have suggested that this motor-related neuronal modulation is not confined to a specific area but pervasive throughout cortical and subcortical brain regions^4–8^. Furthermore, in animals engaged in cognitive tasks, movements that are neither instructed nor necessary for task execution can be aligned to task events and significantly contribute to task-aligned neuronal activity^5–8^. In particular, orofacial movements, i.e., movements of facial parts surrounding the mouth, such as the nose and whiskers, strongly correlate with brain-wide neuronal activities in mice^4, 5, 9^. However, the mechanisms by which the brain generates and coordinates orofacial movements uninstructed but task-aligned remain largely elusive.

Mice well-trained for a stimulus-reward association task frequently exhibit whisker movements immediately following the presentation of a reward-predicting cue^5, 10–13^. The acquisition of liquid rewards through licking also seems to elicit whisker and other facial movements^5, 10–13^. These orofacial movements are often regarded as part of facial expressions related to reward expectation and acquisition^12, 14, 15^. In fact, animals’ behavioral states during task performance can be deduced from the analysis of whisker movements^12^. However, given that licking behavior is phase-locked to breathing, sniffing, and other orofacial activities^16–18^, it remains equivocal whether such orofacial movements are simply concurrent with goal-directed actions or are instead driven by independent neural mechanisms. Moreover, nothing is currently known about which neuronal populations are directly involved in forming task-aligned, uninstructed orofacial movements.

Dopamine (DA) neurons in the ventral tegmental area (VTA) play a central role in mediating motivated behaviors and are supposed to form part of the reward circuit of the brain^19–21^. Transient stimulation of VTA-DA neurons is reinforcing^22–24^ and can replace some aspects of liquid reward^25^. The VTA-DA neurons in mice fire phasically upon reward expectation and acquisition^26–28^, the timings similar to uninstructed whisker movements in reward-based tasks^5, 10, 11^. Therefore, the VTA-DA neurons may be involved in making uninstructed orofacial movements during reward-based tasks.

The primary motor cortex (M1) is another candidate for a brain region that may be involved in uninstructed orofacial movements. There has been compelling evidence that stimulation of specific areas within the orofacial M1 of rodents triggers movements in the whisker and nose^29–33^. Neurons in the orofacial M1 encode various aspects of movements in the whisker and nose^34–36^. Nevertheless, it is unknown whether the orofacial M1 plays a causal role in constituting uninstructed orofacial movements during reward-based tasks.

In this study, we show that distinct whisker movements are induced upon reward predicting cue presentation and reward-acquiring reaction in stimulus-water reward association tasks. Using optogenetic stimulation of VTA-DA neurons (oDAS), we demonstrate that these two types of orofacial movements aligned to the task events could also be elicited in the absence of goal-directed action. Our perturbational analyses revealed that the mesolimbic DA pathway is essential in driving oDAS-aligned orofacial movements but not cue-locked ones, while wM1 is a critical circuit node for driving these two types of movements apart from goal-directed behaviors. Based on these findings, we propose a neural circuit model where neuronal signals converge into wM1 for facilitating uninstructed orofacial movements during a reward-based learning task.

## RESULTS

### Whisker movements during reward-based learning tasks

When mice learn a simple stimulus-reward association task, some orofacial movements become aligned to task events^5, 10, 11^. We first analyzed the whisker movement data in our previous recordings using mice performing a whisker detection task, in which thirsty mice learned to lick for water reward in response to a whisker deflection^37^ (Figure 1A). Expert mice well-trained for the task exhibited a rapid whisker protraction immediately after the reward-associated, brief (1 ms) whisker stimulation in hit trials (Figure 1B), which had a significantly larger amplitude compared to miss trials (hit 15.8 ± 2.2 deg, miss 5.6 ± 1.0 deg, n = 6 mice, p = 0.013; Figure 1C). Such a rapid cue-locked whisker protraction in hit trials was absent in novice mice with low task proficiency (Figure 1B), suggesting that the cue locked whisker protraction becomes evident after learning the task. These expert mice also showed active whisker movements after reward acquisition in hit trials, some of which appeared similar to exploratory whisking^30–32^ (Figure 1B). Total whisker movements (ΣWM) during 1 s after the reward time window were larger in hit trials than in miss trials (hit 381 ± 59 deg, miss 156 ± 31 deg, n = 6 mice, p = 0.0052, Bonferroni’s multiple comparison test; Figure 1C). In contrast to the learning-dependent nature of the cue-locked whisker movements, task learning did not change the ΣWM after hits (novice hit 444 ± 96 deg, n = 6 mice, p = 0.93, Bonferroni’s multiple comparison test vs. expert hit; Figure 1C).

**Figure 1.**
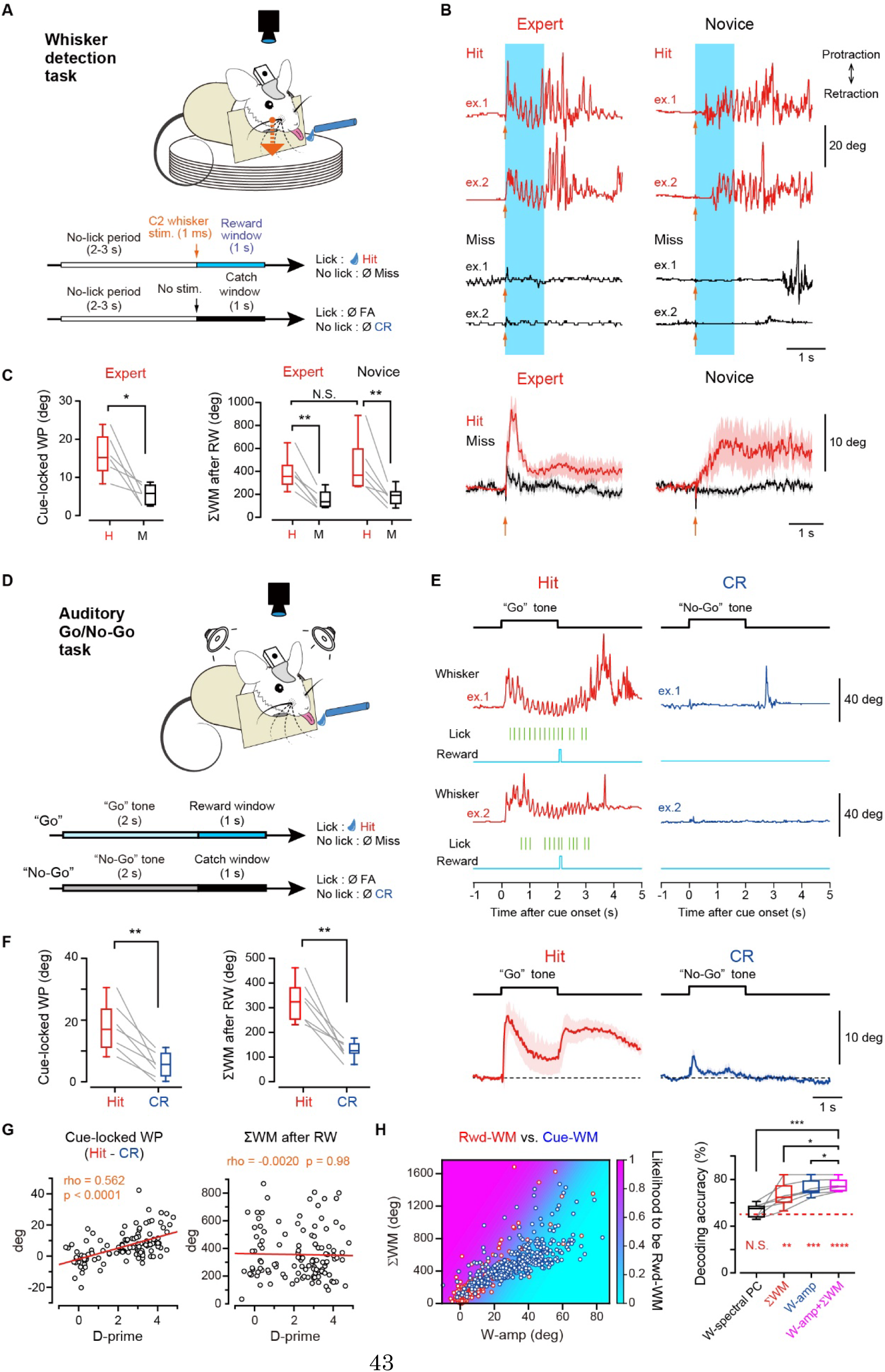
Stereotypical whisker movements during stimulus-reward association tasks. (A) Schematic for the whisker detection task. FA: false alarm, CR: correct rejection. (B) Top, example traces of the whisker position in hit (red) and miss (black) trials during the whisker detection task, obtained from an expert (left) and novice (right) mouse. Arrow: whisker stimulation. Light blue shadow: the reward time window (RW). Bottom, grand average traces of the whisker position (*n* = 6 mice for each). Shadows indicate ± SEM. (C) Amplitudes of the cue-locked whisker protraction (WP) (left) and the total whisker movements (ΣWM) 0–1s after RW (right), in hit (H) and miss (M) trials during the whisker detection task (n = 6 mice for each). ** p < 0.01, *p < 0.05, N.S., not significant, paired *t* test (cue-locked WP) and Bonferroni’s multiple comparison test (ΣWM after RA) (D) Schematic for the auditory Go/No-Go task. (E) Example (top) and grand average (bottom, n = 7 expert mice for each) traces of the whisker position in hit (red) and CR (blue) trials, and other task variables during the auditory Go/No-Go task. (F) Cue-locked WP amplitude (left) and ΣWM 0–1s after RW (right) during the auditory Go/No-Go task (n = 7 expert mice). **p < 0.01, paired *t* test. (G) Left, the difference in the cue-locked WP amplitude between hit and CR trials (left) and ΣWM 0–1s after RW of hit trials (right) as a function of task proficiency (d-prime), obtained from 10 mice. Red: linear regression line. (H) Left, an example decoding analysis of whisker movements at 0–1 s after the cue onset (Cue-WM, blue) and 0–1 s after RW (Rwd-WM, red), obtained from an expert mouse. Right, decoding accuracies by the models for discrimination between Cue-WM and Rwd WM (see STAR★Methods) (n = 7 for each). The dashed red line indicates the chance level. ****p < 0.0001, ***p < 0.001, ** p < 0.01, *p < 0.05, N.S., not significant, Dunnett’s multiple comparison test vs. WP+ΣWM (shown in black) or one-sample *t* test vs. chance level (shown in red). Thin lines in (C), (F), and (H) and open circles in (G) and (H) indicate individual data. See also Figure S1.

We next examined whether the task-aligned whisker movements seen in the whisker detection task are specific for a task involving whisker sensation. We trained mice for an auditory Go/No-Go task where whisker sensation or motion is not required for task execution. We set the reward time window (1 s) after a 2-s auditory cue presentation (Figure 1D) to well separate the timings of expectation and acquisition of water reward. Expert mice for the auditory task exhibited a rapid cue-locked whisker protraction (17.4 ± 3.0 deg, n = 7 mice) and active whisker movements after reward acquisition (ΣWM 0–1 s after the reward window, 320 ± 32 deg, n = 7 mice) in hit trials, both of which were more prominent than in correct rejection (CR) trials (Figures 1E and 1F). In the auditory task, both Go and No-Go sound cues equally elicited a rapid whisker protraction in novice mice (Figures S1A and S1B), which might be a reflex motor action to a salient auditory stimulus. However, the difference in the amplitude of the cue-locked whisker protraction between hits and CR trials became more prominent as the mice learned the task (Figure 1G). In contrast, task learning did not change ΣWM after hits (Figure 1G). These results are essentially the same as those in the whisker detection task. Our machine-learning models accurately discriminated between whisker movements during hit trials and those during CR trials in expert mice (Figure S1C), indicating that the movements of a single whisker can provide information about the behavioral states of mice. Additionally, our models could distinguish between cue-locked whisker movements and those that occurred after reward acquisition, with a decoding accuracy of 75.0 ± 2.0% (by a model that uses ΣWM and WP amplitude, n = 7 mice, *p* < 0.0001, one sample *t*-test against the chance level [50%]; Figure 1H). Thus, whisker movements with distinct properties are observed at the timings of reward expectation and a second after reward acquisition in stimulus-reward association tasks.

### Orofacial movements evoked by oDAS

The uninstructed whisker movements seen in our reward-based tasks (Figure 1) may be associated with anticipatory or reward-acquiring licking which has a specific pattern frequency (5–8 Hz) and is phase-locked to whisker movements^16–18^ (Figures S1D–S1F). We next built an experimental setting to reward mice without inducing any motor actions required for task execution. We implanted an optic fiber unilaterally over the VTA of transgenic mice expressing channelrhodopsin 2 (ChR2) in DA neurons (DAT-ChR2, Figure 2A). We first tested whether a brief (1 s) train of optogenetic stimulation of the VTA-DA neurons (named oDAS), which is known to be rewarding and reinforcing^22–25^, can induce facial movements. An oDAS with 20 pulses at 20 Hz elicited facial movements at the rostral parts including whiskers and nose, but not around ears and eyes, in awake head-restrained mice (Figure 2B). We therefore further analyzed oDAS-induced whisker and nose movements by monitoring the frontal face of mice from a top view (Figures 2C–2J). Our strongest oDAS with 20 pulses at 20 Hz induced rhythmic whisking with a high probability (83.8 ± 8.5 %, n = 7 mice; Figure 2F). The average whisker angle during oDAS was protracted by 11.6 ± 2.6 degrees (n = 7 mice, 20 pulses at 20 Hz; Figure 2F). The probability of whisking initiation, ΣWM during oDAS, and the average and maximum amplitude of whisker movements all increased, depending on the stimulus frequency (Figures 2D–2F) and photo-stimulus intensity (Figures S2A and S2B). In contrast, in control Ai32 mice without ChR2-expression, photostimulation of VTA did not evoke any consistent whisker movements (Figures 2D–2F, S2A and S2B). Such protracted rhythmic whisking evoked by oDAS resembled whisker movements upon reward acquisition in reward-based learning tasks (Figures 1B and 1F). The oDAS-induced whisking had an average latency of 402 ± 51 ms (n = 7 mice, with 20 pulses at 20 Hz; Figure 2J).

**Figure 2.**
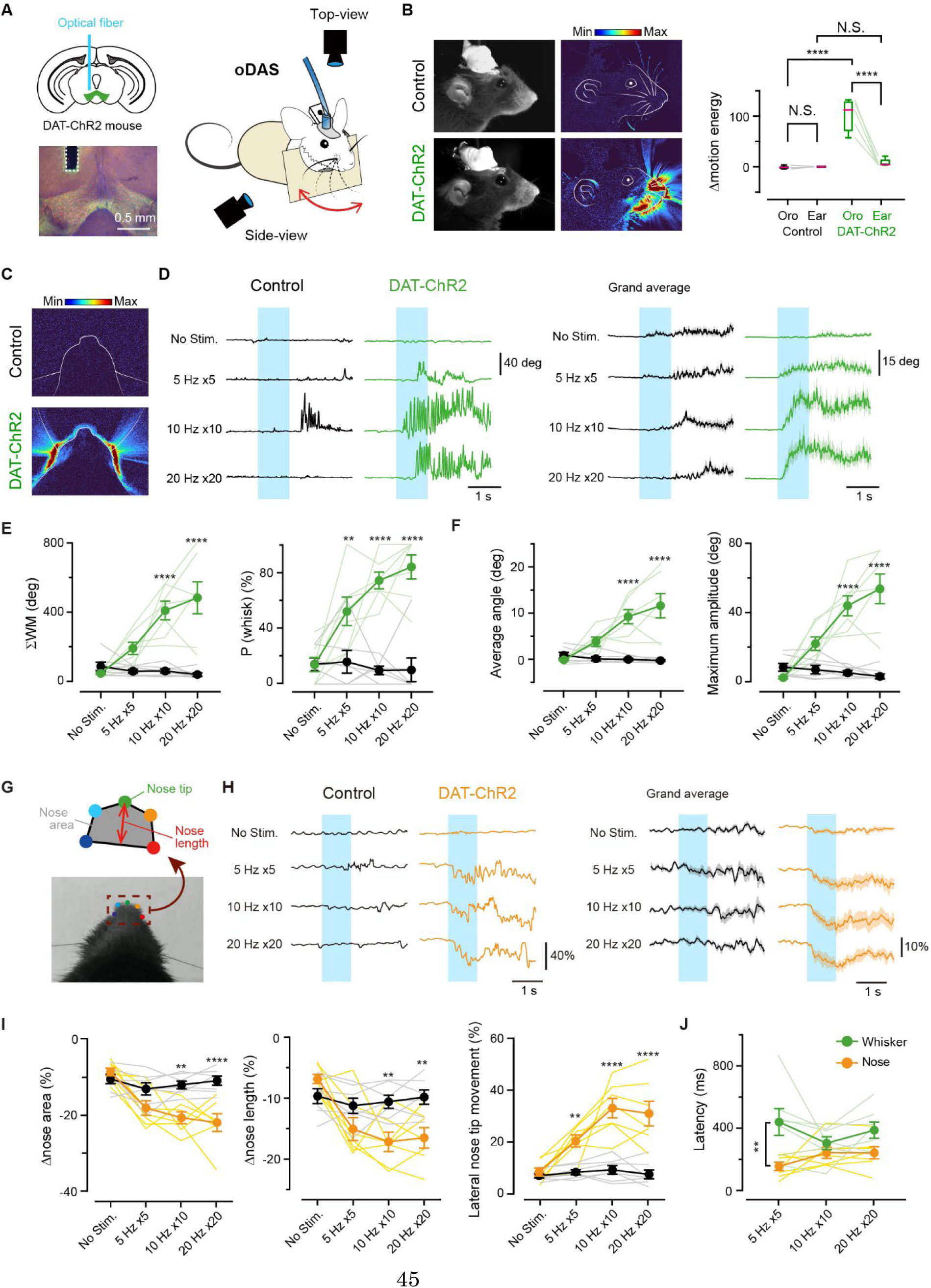
Orofacial movements induced by transient activation of VTA-DA neurons. (A) Left, schematic for the optical fiber location for the oDAS experiments (top), and an epifluorescence image of a coronal section containing VTA and the trace of the inserted fiber (dashed line) (bottom). Green: ChR2-eYFP, red: tyrosine hydroxylase, blue: DAPI. Rignt, schematic for the oDAS experiment. (B) Left, single example video frames from individual mice. Middle, motion energy heat maps overlayed onto the face line drawings from the mice shown in Left. Right, quantifications for motion energy around the orofacial part (Oro) and the ear during oDAS. Medians of the box plots are shown in red. (C) Example motion energy heat-maps overlayed onto the top-view face line. (D) Left, example traces of the whisker position upon oDAS (light blue shadow) with different stimulus frequencies, obtained from a control (black) and DAT-ChR2 (green) mouse. Right, grand average of the whisker position upon oDAS (control: n = 7 mice; DAT-ChR2: n = 7 mice). Shadows of traces: ± SEM. (E) ΣWM (left) and P(whisk) (the probability of whisking initiation, right) during oDAS. (F) Average (left) and maximal (right) amplitude of whisker protraction (WP) angle during oDAS. (G) Schematic for the nose analysis (top) and a top-view snapshot of mouse face (bottom). (H) Same as (B) but for traces of the top-view nose area from control (black) or DAT-ChR2 (orange) mice. (I) Quantifications for the changes in nose area (left), nose length in the anterior-posterior axis (middle), and lateral movements of the nose tip (right). (J) Latencies of movement of the whisker (green) and nose (orange) upon oDAS. Filled circles and error bars show mean ± SEM. Lightly colored lines correspond to individual data. ****p < 0.0001, **p < 0.01, N.S., not significant, Bonferroni’s multiple comparison test, control vs. DAT-ChR2 in (E), (F), and (I) or whisker vs. nose in (J). See also Figure S2.

We next analyzed nose movements evoked by oDAS using a deep learning-based toolbox DeepLabCut^38^ (Figure 2G). Mice exhibited nose twitches upon oDAS: the top-view nose area decreased (peak change from baseline: 22.0 ± 2.3%, n = 7) (Figures 2H and 2I), and the nose length in an anterior-posterior axis became shortened (peak change from baseline: 16.5 ± 1.6%, n = 7) (Figure 2I), indicating contractions of the nasal muscle. The oDAS also induced a lateral movement of the nose tip (peak lateral movement as the percentage of the mean baseline nose length: 31.0 ± 4.8%, n = 7) (Figure 2I). The magnitude of such nose movements depended on the stimulus frequency (Figure 2H and 2I) and photo-stimulus intensity (Figures S2A and S2C). Upon oDAS stimulation, nose and whisker movements were highly correlated (Figure S2D). The latencies of nose movement initiation and whisking initiation were similar, except for when using the weakest stimulation (Figure 2J). Thus, transient activation of VTA-DA neurons is sufficient to facilitate movements in the whisker and nose. In contrast, optogenetic stimulation of DA neurons in the substantia nigra pars compacta (SNc), which are implicated in the coordination of locomotion in mice^39, 40^, did not induce whisker and nose movements (Figures S2E–S2G).

### Orofacial movements time-locked to reward-predicting cues

In our liquid reward-based tasks, the cue-locked whisker protraction was immediately followed by anticipatory or goal-directed licking (Figures 1 and S1D–S1F). We next examined whether the cue-locked whisker protraction can be induced without anticipatory licking. We presented a sound cue (5 s) paired with oDAS during the last 1 s of the cue to head-restrained DAT-ChR2 or control (Ai32) mice (Figures 3A and 3B). Mice experienced ∼20 trials of the paired stimulation per day, with random intertrial intervals between 180 and 240 s. On the first day of this conditioning (day 1), both DAT-ChR2 and control mice similarly exhibited a whisker protraction immediately after the sound presentation with a high probability (Figure 3C), which might represent a reflex motor action to a salient auditory stimulus as observed in the auditory Go/No-Go task (Figures S1A and S1B). On day 2, however, DAT-ChR2 mice exhibited a rapid, cue-locked whisker protraction (24.3 ± 1.6 deg, n = 26 mice) (Figure 3B) with a significantly larger magnitude than that on day 1 (18.3 ± 1.3 deg, n = 26 mice, p < 0.0001, Bonferroni’s multiple comparison test; Figures 3C and S3A). The magnitude of the cue-locked whisker protraction continued to increase over four days of learning (Figure S3B). In contrast, in control mice, the magnitude and probability of the cue-locked whisker protraction significantly decreased on day 2 (Figures 3B, 3C, and S3A). These results indicate that learning the sound-oDAS pairing can elicit cue-locked whisker protraction even in the absence of goal-directed action. The latency of whisker protraction after the onset of the sound presentation was 98 ± 7 ms (n = 26 mice) on day 2 (Figure 3G), which was significantly shorter than that of oDAS-induced whisker motion (p < 0.0001, with 20 pulses at 20 Hz, Mann-Whitney *U* test, Figure 2J). In addition to the whisker protraction, a quick nose twitch was also elicited immediately after the sound onset in DAT-ChR2 mice on day 2 (peak change in the nose area: 25.1 ± 2.2%, n = 26 mice; peak change in the nose length: 22.2 ± 2.0%, n = 26 mice; peak lateral movement of the nose tip: 36.1 ± 2.2%, n = 26 mice), which was absent in control mice without ChR expression on day2 (Figures 3D– 3F). The latency of the cue-locked nose twitch (156 ± 10 ms, n = 26 mice) was significantly longer than that of the cue-locked whisker protraction (p = 0.0008, two-way ANOVA; Figure 3G) but still shorter than that of oDAS-induced nose movement (257 ± 38 ms, upon oDAS at 20 Hz for 1 s, n = 7 mice, p = 0.0006, unpaired *t* test). The amplitudes of the nose twitch and whisker protraction at the cue onset were weakly but significantly correlated (Figure S3C). Control mice showed small reflex movements in the whisker and nose at the cue onset in a small fraction of the trials on day 1 (Figures 3C, 3E, 3F, and S3A). Its magnitude and probability significantly decreased with training (Figures 3C, 3E, 3F, and S3A), suggesting that learning of sound-oDAS association should cause the cue-locked orofacial motions in DAT-ChR2 mice on day 2. In this sound-oDAS association task, we also observed oDAS-aligned whisker movements (Figure 3B). The magnitude of whisker movements and the probability of whisking during oDAS in this task did not change over five days, with a slight tendency to be attenuated after three days, although not statistically significant (Figure S3D).

**Figure 3.**
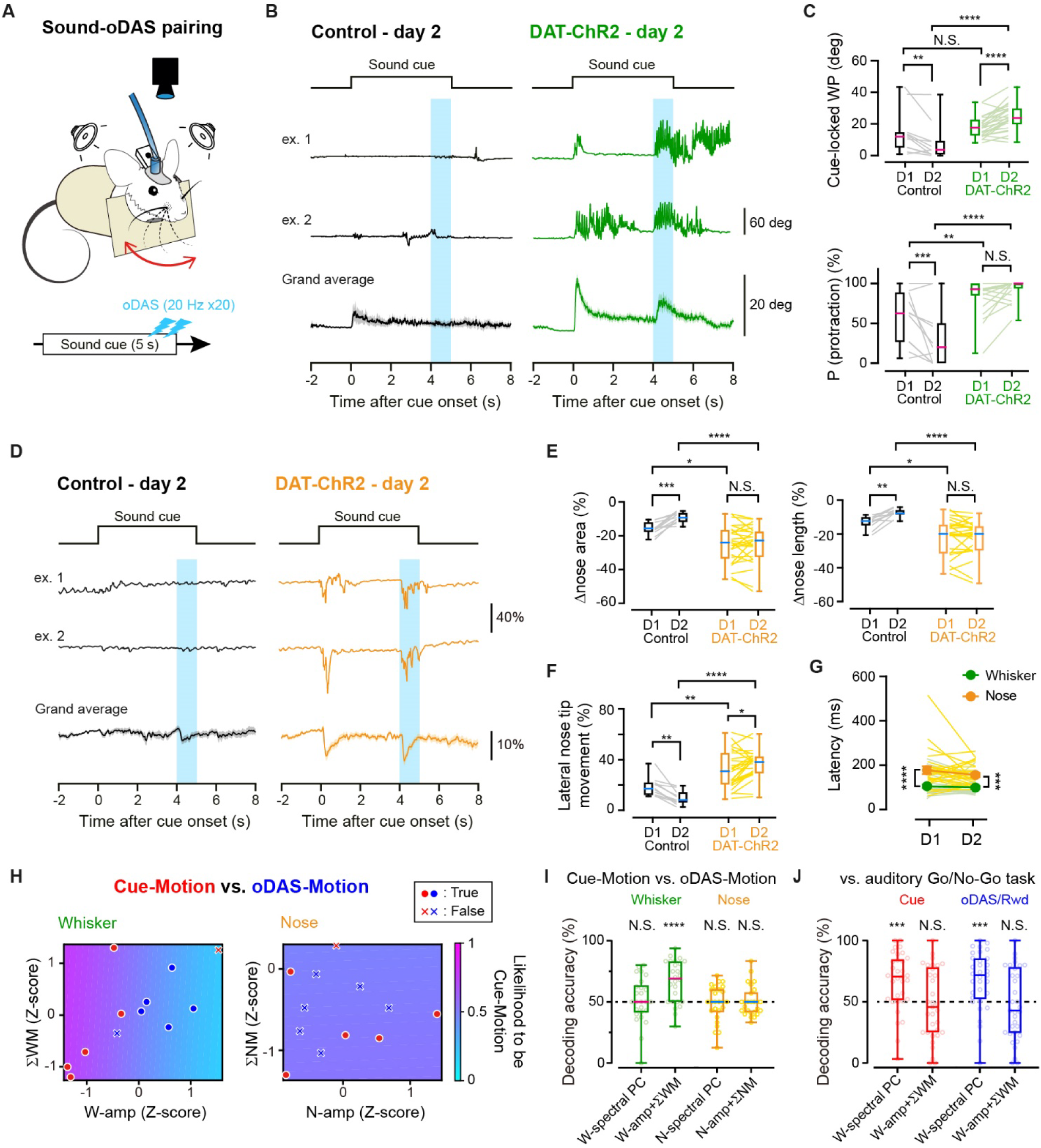
Orofacial movements during a stimulus-oDAS association task. (A) Schematic for the sound-oDAS pairing conditioning. (B) Example and grand average traces of the whisker position during the sound-oDAS pairing conditioning on day 2, obtained from a control (black) and DAT-ChR2 (green) mouse (control: n = 11 mice; DAT-ChR2: n = 26 mice). The timings of the sound cue presentation (black line) and oDAS (light blue shadow) are indicated. Shadows of grand average traces: ± SEM. (C) The amplitude of cue-locked whisker protraction (WP, top) and P(protraction) (protraction probability, bottom) on day (D) 1 and 2. (D) Same as (B) but for traces of the top-view nose area from control (black) or DAT-ChR2 (orange) mice. (E) Quantifications for the changes in the nose area and nose length (F) Quantifications for the changes in lateral movements of the nose tip. (G) Latencies of cue-locked whisker protraction (green) and nose tip movements (orange) upon sound cue presentation. (H) Example decoding analyses of whisker (left) and nose (right) movements at 0–1 s after the cue onset (red, Cue-Motion) and during 1-s oDAS (blue, oDAS-Motion), obtained from an expert mouse (see STAR★Methods). True: correctly predicted; False: incorrectly predicted. (I) Decoding accuracies by the models for discrimination between cue-locked and oDAS aligned whisker/nose movements during the sound-oDAS task (*n* = 26 mice). The dotted line indicates the chance level. (J) Same as (I) but for discrimination between whisker motions during the sound-oDAS task and those during the auditory Go/No-Go task. Medians of the box plots are shown in magenta or cyan. Filled circles and error bars in (G) show mean ± SEM. Lightly colored lines correspond to individual data. ****p < 0.0001, ***p < 0.001, **p < 0.01, N.S., not significant, two-way ANOVA followed by Bonferroni’s multiple comparison test, except for one-sample *t* test vs. the chance level in (I). See also Figure S3.

Our machine-learning models, utilizing ΣWM and WP amplitude, could distinguish oDAS-induced and cue-locked whisker movements (decoding accuracy, 66.6 ± 3.4%, n = 26, p < 0.0001, one-sample *t* test against the chance level [50%]; Figures 3H and 3I). This suggests that the reward-predicting cue and oDAS induce whisker movements with distinct properties. However, our models based on nose movements were unable to predict the timing of the data (cue onset or during oDAS) (Figures 3H and 3I). Additionally, we compared whisker movements during the sound-oDAS and auditory Go/No-Go tasks. The models based on ΣWM and WP amplitude could not distinguish cue-locked or oDAS/reward-aligned whisker movements induced in these two types of tasks (Figure 3J). However, the models based on spectral PC could discriminate between them (Figure 3J). The spectral PC primarily derives from oscillatory whisker movements associated with goal-directed licking (Figures S1C–S1F). These results suggest that whisker movements during the sound-oDAS pairing task are indistinguishable from those during water-reward-based learning tasks, except for the presence of licking-associated oscillatory movements.

### Involvement of accumbal D1R in driving orofacial movements

We next investigated the neuronal circuits downstream of VTA DA neurons for inducing reward-related orofacial movements. Stimulation of VTA DA neurons is known to activate medium spiny neurons expressing D1 receptors (D1Rs) in the nucleus accumbens (NAc) through the mesolimbic DA pathway^41^. We therefore examined the role of D1Rs in reward related orofacial movements. Intraperitoneal (IP) injection of a D1 receptor (D1R) antagonist, SCH23390 (0.3 mg/kg), abolished the oDAS-induced whisking (ΣWM: saline = 487 ± 127 deg [n = 7 mice], SCH23390 = 32.4 ± 5.3 deg [n = 7 mice], p = 0.013, paired *t* test; whisking probability: saline = 81.6 ± 11.2% [n = 7 mice], SCH23390 = 5.4 ± 5.4% [n = 7 mice], p = 0.031, Wilcoxon signed rank test) and nose twitches (peak change in the nose area: saline = 25.0 ± 5.5% [n = 7 mice], SCH23390 = 5.0 ± 1.3% [n = 7 mice], p = 0.011, paired *t* test) in DAT-ChR2 mice (Figures 4A, 4B and S4A), suggesting the involvement of D1Rs. The IP injection of SCH23390 also reduced the spontaneous whisker movements (ΣWM per a second: saline = 159 ± 25 deg/s [n = 9 mice], SCH23390 = 55.2 ± 8.1 deg/s [n = 9 mice], p = 0.0058, paired *t* test; Figure 4A), suggesting a general sedative effect of the D1R antagonist. To specifically examine the role of D1Rs in the NAc in driving oDAS-induced orofacial movements, we used CRISPR-Cas9-mediated *in vivo* genome editing. We knocked out D1Rs in the NAc of DAT-ChR2 mice by local injection of an AAV vector harboring saCas9 with guide RNA targeting the *Drd1a* gene (AAV-gRNA), which has been previously shown to have a high cleavage rate^42^ and no off-target effects (Figure 4C, S4B and S4C). These NAc-D1R-KO mice did not show whisker movements or nose twitches upon oDAS (Figures 4C, 4D and S4D). In contrast, control mice injected with a control AAV saCas9-gRNA vector (AAV-Control) exhibited oDAS-induced whisker movements and nose twitches (Figures 4C, 4D and S4D). Spontaneous whisker movements were unaffected in NAc-D1R-KO mice (Figure 4C). These results suggest the involvement of accumbal D1Rs in oDAS-induced orofacial movements. Moreover, optogenetic activation of the axons of VTA-DA neurons in the NAc induced whisker movements and nose twitches (Figures 4E, 4F and S4E). Our results thus indicate that the mesolimbic DA pathway mediates oDAS induced orofacial movements through the activation of D1Rs in the NAc.

**Figure 4.**
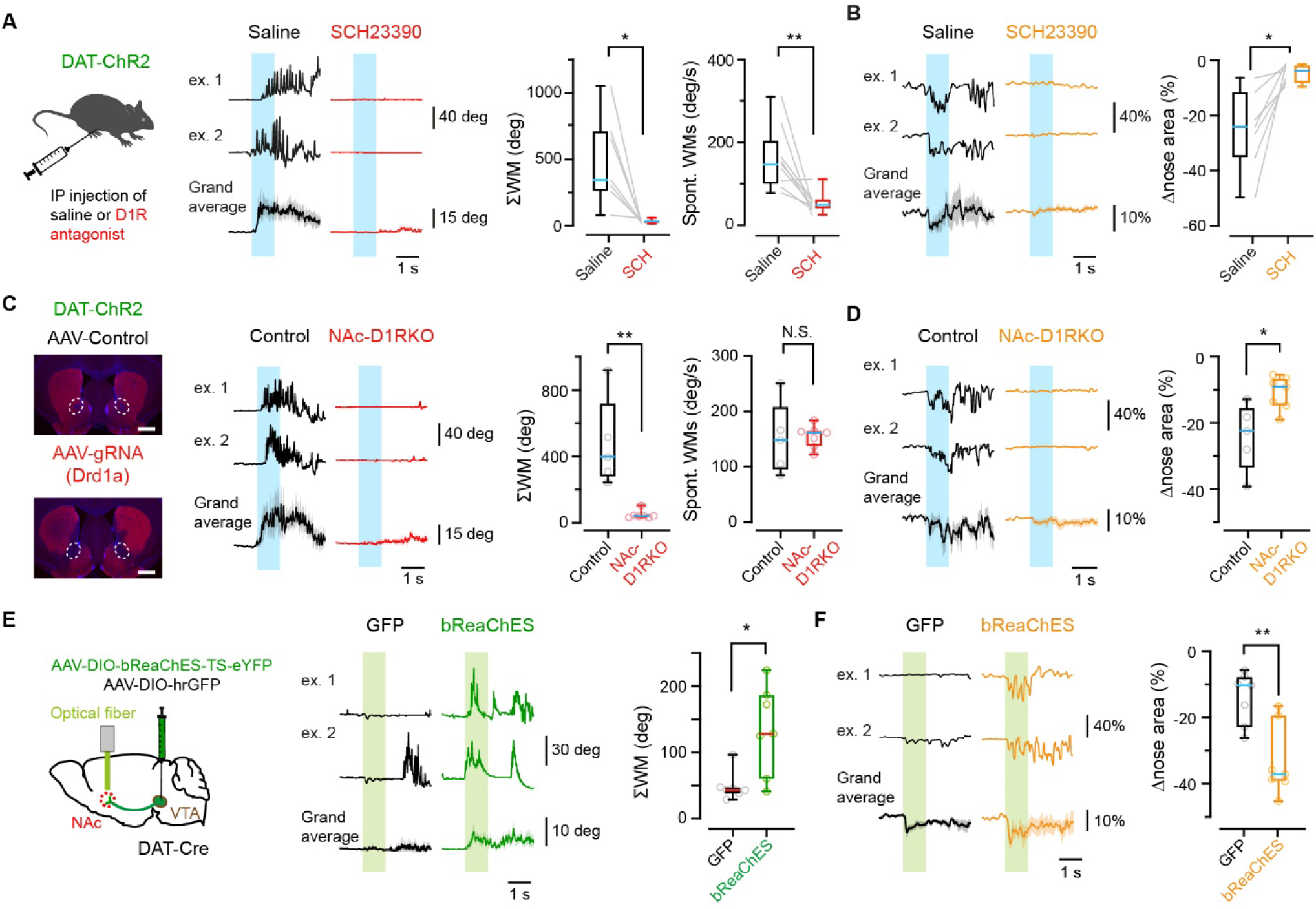
Essential involvement of accumbal D1Rs in oDAS-induced orofacial movements. (A) Left, schematic for the experiment. Middle, example and grand average traces of the whisker position upon oDAS (light blue shadow) in trials with IP injection of SCH23390 (red) and saline (black) (n = 7 mice). Right, ΣWM during oDAS and the magnitude of spontaneous whisker movements (Spont. WMs, n = 9 mice). (B) Example and grand average traces (left) and quantifications (right) of the top-view nose area in trials with IP injection of saline (black) or SCH23390 (orange) (n = 7 mice). (C) Left, epifluorescence images of coronal sections containing NAc (dashed circles) from mice with bilateral injection of AAV with guide RNA targeting *Drd1a* (bottom) or control AAV with shuffled RNA (top). Red, Drd1 immunoreactivity, blue: DAPI, Scale bar: 1 mm. Middle, example and grand average traces of the whisker position in NAc-D1R-KO (red) and control (black) mice upon oDAS (control: n = 5 mice; NAc-D1R-KO: n = 7 mice). Right, ΣWM and the magnitude of Spont. WMs during oDAS (control: n = 5 mice; NAc D1R-KO: n = 7 mice). (D) Same as (B) but for data from NAc-D1R-KO (orange) and control (black) mice. (E) Left, schematic for the optogenetic experiment. Middle, example and grand average traces of the whisker position upon oDAS in bReaChES (green) and GFP (black) expressing mice. Right, data for each mouse (thin lines) and box plots for the ΣWM during oDAS (n = 7 mice for each). (F) Same as (B) but for data from bReaChES (orange) and GFP (black) expressing mice. Green shadow: stimulation timing. Medians of the box plots are shown in magenta or cyan. Shadows of grand average traces: ± SEM. Open circles and lightly colored lines correspond to individual data. **p < 0.01, *p < 0.05, N.S.: not significant, paired *t* test in (A) and (B), Mann-Whitney *U* test (ΣWM in (C), and (E)) or unpaired *t* test (Spont. WMs in (C), and (D)). See also Figure S4.

We further tested whether D1Rs are also involved in generating the cue-locked whisker protraction. We performed an IP injection of saline or SCH23390 in the mice that had learned the sound-oDAS pairing for two days and analyzed orofacial movements upon the paired stimulation on the third day. Even though the IP injection of SCH23390 dramatically reduced spontaneous and oDAS-induced whisker movements (Figure 4A), it did not affect the cue-locked whisker protraction after learning of the sound-oDAS pairing (cue-locked whisker protraction amplitude: saline = 33.8 ± 4.4 deg [n = 6 mice], SCH23390 = 30.0 ± 4.0 deg [n = 6 mice], p = 0.35, paired *t* test) (Figures 5A and 5B). The D1R antagonist did also not significantly attenuate nose twitches time-locked to the sound cue (peak change in the nose area: saline = 24.0 ± 3.7 deg [n = 6 mice], SCH23390 = 18.3 ± 3.4 deg [n = 6 mice], p = 0.20, paired *t* test) (Figures 5C and 5D). Thus, the oDAS-induced and cue-locked orofacial movements are differentially driven by D1R-dependent and independent neuronal mechanisms, respectively. We further tested whether another DA receptor, D2 receptor (D2R), is involved in cue-locked orofacial movements. IP injection of SCH23390 together with a D2R antagonist, raclopride (3 mg/kg), did not affect cue-locked whisker movements (Figures 5E and 5F), but it largely attenuated the cue-locked nose movements (Figures 5G and 5H). These results suggest that D2Rs regulate whisker and nose movements differentially in response to reward-predicting cue presentation.

**Figure 5.**
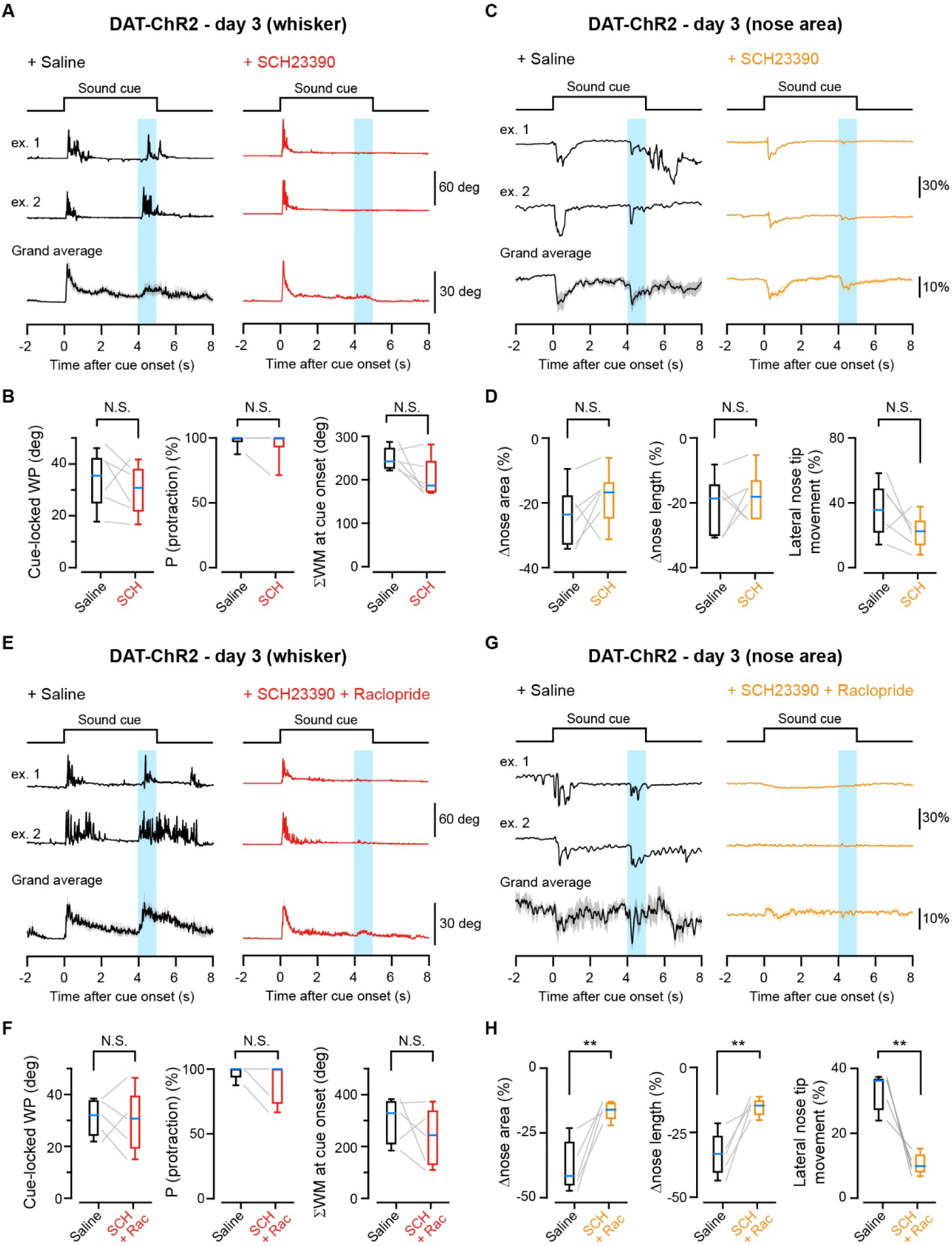
Orofacial movements upon reward expectation do not require D1Rs. (A) Example and grand average traces of the whisker position in trials with SCH23390 or saline-injection in DAT-ChR2 mice during sound-oDAS pairing experiments on day 3 (n = 6 mice). The timings of the sound cue presentation (black line) and oDAS (light blue shadow) are indicated. Shadows of grand average traces: ± SEM. (B) Quantification of cue-locked whisker movements with SCH23390 or saline injection. (C) Same as (A) but for traces of the top-view nose area. (D) Same as (B) but for the parameters of nose movement at the cue onset. (E–H) Same as (A–D) but with D1R- and D2R-antagonists- or saline-injection (n = 5 mice). Medians of the box plots are shown in cyan. Thin lines correspond to individual data. **p < 0.01, N.S.: not significant, paired *t* test for all comparisons in (B), (D), (F), and (H), except for P(protraction) in (B) and (F) with Wilcoxon signed rank test.

### Whisker M1 facilitates uninstructed orofacial movements

Mouse whisker primary motor cortex (wM1) plays a critical role in triggering exploratory whisking^30–32^. Having identified D1R-independent neuronal mechanisms driving the cue locked orofacial movements, we next examined the involvement of wM1 in reward-related orofacial movements using DAT-ChR2 mice. Silencing of wM1 by injection of muscimol (5 mM in Ringer’s solution, total 400 nl), a GABAA-receptor agonist, led to a dramatic drop in the magnitude and probability of oDAS-induced whisker movements (ΣWM: before muscimol = 243 ± 40 deg [n = 6 mice], after muscimol = 55.0 ± 12.0 deg [n = 6 mice], p = 0.0095, paired *t* test; whisking probability: before muscimol = 59.9 ± 6.1% [n = 6 mice], after muscimol = 17.1 ± 6.4% [n = 6 mice], p = 0.0027, paired *t* test) (Figures 6A and 6B). Inactivation of wM1 by muscimol also attenuated oDAS-induced nose twitches (peak change in the nose area: before muscimol = 24.2 ± 1.6% [n = 6 mice], after muscimol = 5.6 ± 1.3% [n = 6 mice], p = 0.0001, paired *t* test; Figures 6C, 6D and S5A). Injection of Ringer’s solution in wM1 did not affect these oDAS-induced orofacial movements (Figures 6C, 6D, and S5A). Thus, neuronal activity in wM1 is required for the generation of oDAS-induced orofacial movements.

**Figure 6.**
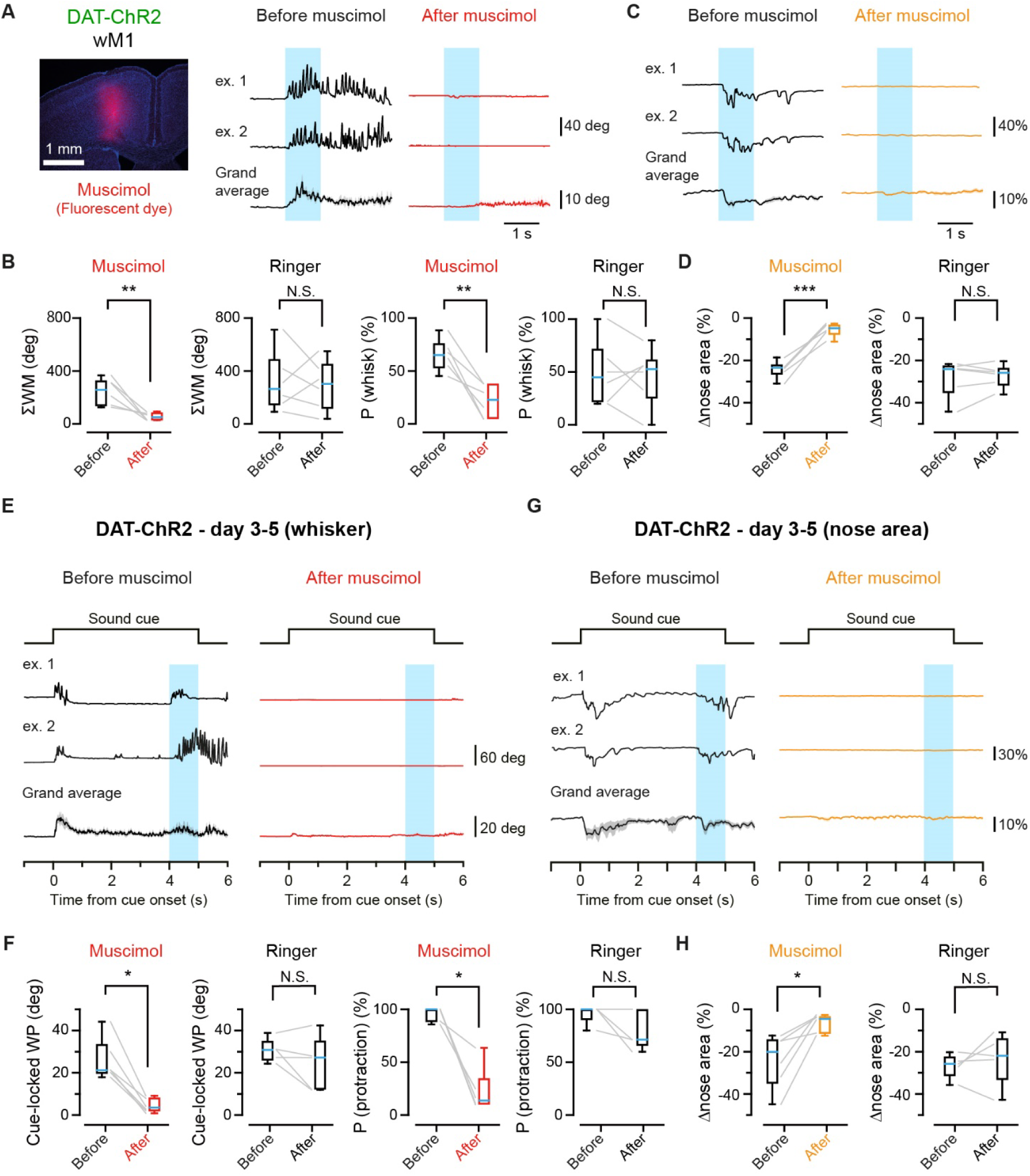
Dominant role of wM1 in driving reward-related orofacial movements. (A) Left, an epifluorescence image of a coronal section indicating the muscimol injection site. A small volume of Chicago sky blue (red) was dissolved into the vehicle. Blue: DAPI Right, example and grand average traces of the whisker position upon oDAS (light blue shadow) before (black) and after (red) muscimol injection in wM1 of DAT-ChR2 mice (n = 6 mice). (B) ΣWM and the probability of whisking initiation in trials before (black) and after (red) injection of muscimol or Ringer’s solution (n = 6 mice for each). (C) Example and grand average traces of the nose area before (black) and after (orange) muscimol injection into wM1. (D) Same as (B) but for the change in the nose area. (E) Example and grand average traces of the whisker position before (black) and after (red) muscimol injection into wM1 of DAT-ChR2 mice during sound-oDAS pairing experiments. (F) Same as (B) but for the amplitude and occurrence probability of cue-locked whisker protraction (WP) during sound-oDAS pairing experiments. (G) and (H) Same as (E) and (F) but for the change in the nose area. Medians of the box plots are shown in cyan. Thin lines correspond to individual data. Shadows of grand average traces: ± SEM. ***p < 0.001, **p < 0.01, *p < 0.05, N.S., not significant, paired *t* test (all of (B), (D) and (H) and cue-locked WP for Ringer in (F)) or Wilcoxon signed rank test (two panels for Muscimol and P(protraction) for Ringer in (F)). See also Figure S5.

We further investigated whether the cue-locked orofacial movements during the sound-oDAS pairing stimuli also depend on wM1 activity. We used DAT-ChR2 mice that learned the sound-oDAS pairing for 2–4 days and tested injection of muscimol or Ringer’s solution into wM1 on the next day. Inactivation of wM1 by muscimol largely attenuated the cue-locked whisker protraction (amplitude: before muscimol = 25.8 ± 4.0 deg [n = 6 mice], after muscimol = 4.6 ± 1.4 deg [n = 6 mice], p = 0.031, Wilcoxon signed rank test; probability: before muscimol = 95.8 ± 2.7% [n = 6 mice], after muscimol = 22.7 ± 8.5% [n = 6 mice], p = 0.031, Wilcoxon signed rank test; Figures 6E, 6F, and S5B) and nose twitch (peak change in the nose area: before muscimol = 24.0 ± 5.2% [n = 6 mice], after muscimol = 6.5 ± 1.7% [n = 6 mice], p = 0.013, paired *t* test; Figure 6G, 6H, and S5C). Injection of Ringer’s solution did not affect the cue-locked orofacial movements (Figure 6G, 6H, and S5C). Our results thus suggest that wM1 activity is crucial for the generation of both oDAS-induced and cue locked orofacial movements.

### Representation of uninstructed orofacial movements in wM1

wM1 thus appears to have an essential role in driving both oDAS-induced and cue-locked orofacial movements. Therefore, it is of interest to investigate the neuronal signals in wM1 during uninstructed orofacial movements. Using multisite silicone probes, we measured the action potential firing in wM1 of head-restrained DAT-ChR2 mice during stimulus-oDAS pairing stimulation, where both oDAS-induced and cue-locked orofacial movements could be induced in single trials. We performed recordings in four DAT-ChR2 mice that had already experienced the stimulus-oDAS pairing conditioning for three days. The reward-related orofacial behaviors (the amplitude and probability of cue-locked whisker protraction and the magnitude and probability of oDAS-aligned whisker movements) of these mice on the recording day were essentially the same as those on day 2 (Figures S6A and S6B). Among the 174 units we identified in wM1, 103 units (59.2 %) significantly increased or decreased their firing rate at the cue onset and/or during oDAS. To examine the correlation between the activity of individual neurons and whisker behavior, we first excluded trials where spontaneous whisking started during the pre-trial period (1 s) from analysis and then categorized trials based on the behavior (Figures 7A and 7B). In 20.8% of trials, both cue locked whisker protraction and whisking during oDAS were observed (CL+OA trials, n = 4 mice). There were also trials with only cue-locked whisker protraction but not oDAS-aligned whisker movements (CL trials, 37.6%) and those with only oDAS-aligned whisker movements but not cue-locked whisker protraction (OA trials, 11.9%). We analyzed a correlation between neuronal activities and cue-locked whisker protraction amplitude in CL+OA and CL trials or variance of whisker position during oDAS in CL+OA and OA trials. We found the units specifically representing cue-locked whisker protractions (CL cells) with their firing rate increased at the cue onset only when mice showed whisker protractions, while their firing rate did not correlate with oDAS-aligned whisker movements (Figures 7C and S6C). In contrast, those representing oDAS-aligned whisker movements (OA cells) responded to oDAS only when mice showed oDAS-aligned whisker movements while showing no correlation with cue-locked whisker protractions (Figure 7D and S6C). There was also a population of wM1 units representing both cue-locked whisker protractions and oDAS-aligned whisker movements (CL+OA cells; Figure 7E and S6C). In our recording, 12.6% of the units were defined as CL cells, 12.1% as OA cells, and 3.4% as CL+OA cells (Figure 7E). We also analyzed the correlation between the cellular activities in wM1 and nose movements (nose area). We found that only 2.3 % of wM1 cells are significantly correlated only with the nose (but not whisker) movement. In contrast, wM1 contained 20.1 % of cells whose firings are correlated specifically with whisker motion (either cue locked or oDAS-aligned or both of them) and not with nose motion (Figures S6D and S6E). These results suggest that the wM1 region we studied is more related to whisker movements than nose movements.

**Figure 7.**
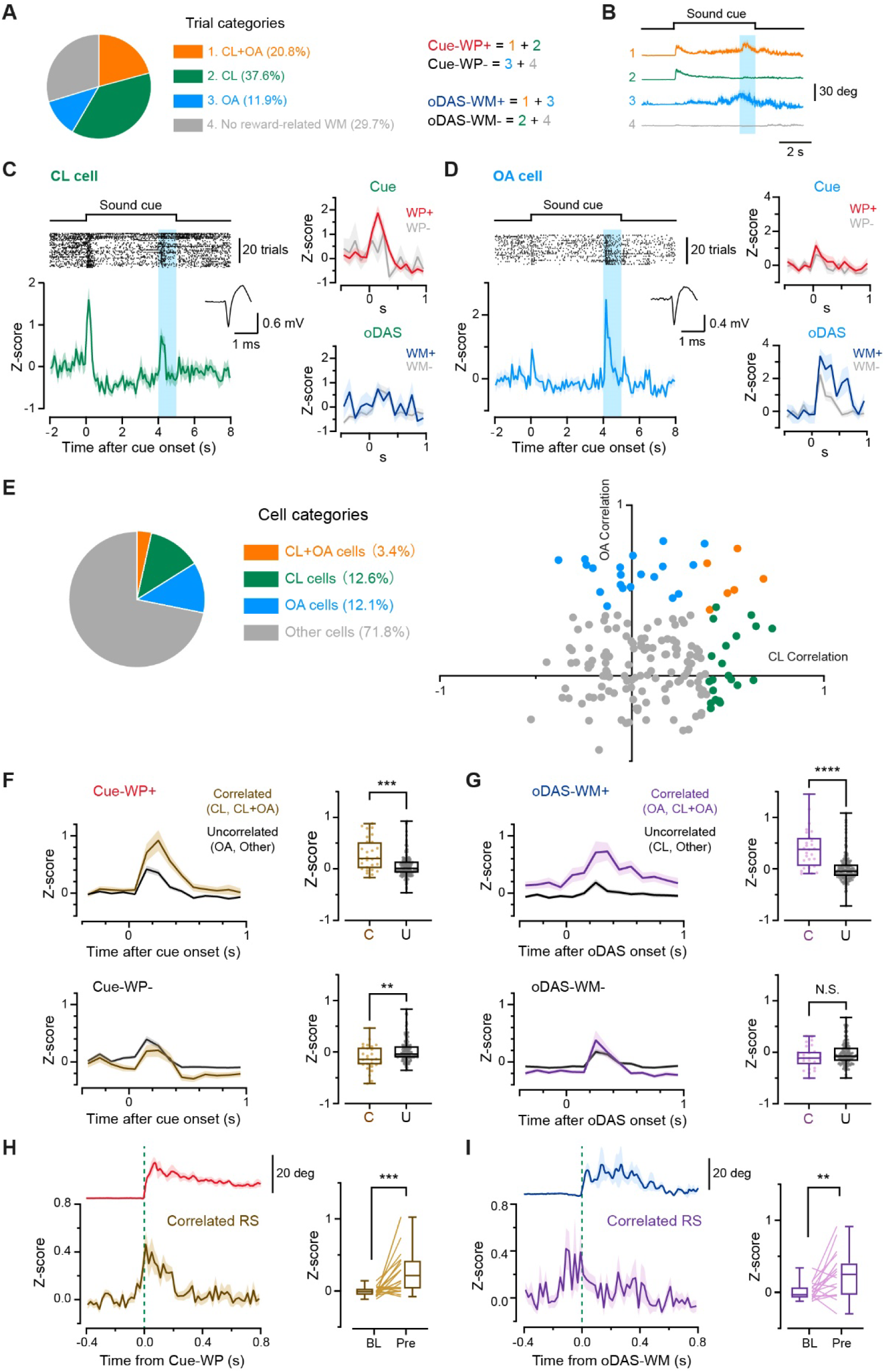
Neuronal representation of cue-locked and oDAS-induced whisker movements in wM1. (A) Summary of the trial categories. CL: trials with cue-locked whisker protraction (Cue-WP); OA: trials with whisker movements during oDAS (oDAS-WM); CL+OA: trials with both Cue-WP and oDAS-WM; +: with motion; -: without motion. (B) Grand average traces of the whisker position in trial categories indicated in (A). (C) Left, example raster plot (top) and corresponding Z-scored peri-stimulus time histogram (PSTH, bottom) obtained from a representative CL cell. The averaged action potential waveform is shown in the inset. Right, Z-scored PSTH of the same cell at around the cue onset (top) and upon oDAS (bottom), comparing data with (+) or without (-) Cue-WP or oDAS-WM. (D) Same as (C) but for a representative OA cell. (E) Left, summary of the cell categories. Right, scatter plot containing 174 neurons from four mice experiencing the sound-oDAS pairing stimulation on the fourth day of conditioning, plotted on the basis of their correlation to the ΣWM during oDAS (OA correlation) and the Cue-WP amplitude (CL correlation). Subsets of neurons showing significant, positive correlation are colored in orange (CL+OA cell), green (CL cell) or cyan (OA cell). (F) Left, average Z-scored PSTHs of Cue-WP-correlated (C) and -uncorrelated (U) cells at around the cue onset, comparing data with (+, top) or without (-, bottom) Cue-WP. Right, average Z-scored firing rates at 0–1 s after the cue onset. (G) Same as (F) but for oDAS-WM-correlated (C) and -uncorrelated (U) cells upon oDAS. (H) Left, Z-scored average PSTHs of Cue-WP-correlated RS cells (*n* = 24 cells) and corresponding grand-averaged whisker positions (n = 4 mice) aligned at the motion onset. Right, averaged Z-score of firing rates at baseline (BL, 0.2–0.4 s before the onset) and pre-motion (Pre, 0.04–0 s before the onset). (I) Same as (H) but for oDAS-WM-correlated RS cells (*n* = 21 cells, 4 mice) upon oDAS. A thick line and shadows indicate mean ± SEM. \*\*\*\**p* < 0.0001, \*\*\**p* < 0.001, \*\**p* < 0.01, N.S., not significant, Mann-Whitney *U* test in (F) and (G) or Wilcoxon signed-rank test in (H) and (I). See also Figures S6 and S7.

In wM1, these cells with correlated firings to each type of whisker motion increased firing rates during the corresponding motion type than uncorrelated cells but not in trials without movements (Figures 7F and 7G). We further separated units into regular spiking (RS) and fast spiking (FS) units, according to their trough-to-peak time of average spike waveform (Figure S7A). We found no apparent specificity in the fraction of FS cells in these CL and OA cells (Figures S7B and S7C) and other stimulus-modified neurons (Figure S7D). The RS cells correlated with cue-locked or oDAS-aligned whisker movements showed an increase in spike rates slightly before or at around the motion onset (Figures 7H and 7I). Our results thus suggest that wM1 contains specific neuronal populations that facilitate the initiation of stereotyped uninstructed whisker movements at the timings of reward expectation and acquisition.

### Stimulation of wM1 triggers whisker and nose movements

Electrical or optogenetic stimulation of wM1 is known to trigger exploratory whisking^30–32^. Our perturbational and electrophysiological studies further indicate that wM1 plays a crucial role as a circuit node in initiating uninstructed whisker and nose movements during sound oDAS pairing stimulation. If this hypothesis is true, stimulating wM1 should result in orofacial movements with a shorter onset latency than those observed after sound presentation or oDAS. Therefore, we investigated the onset latency with which stimulation of wM1 could induce orofacial movements. To express ChR2 in wM1 neurons, we administered AAV (AAV CaMKII-ChR2-mCherry) injections in wM1 (Figure 8A). After 4–5 weeks of expression, we delivered blue light pulses using the same protocol as our strongest oDAS (5 ms, 20 times at 20 Hz) to the wM1 of awake, head-restrained mice (Figure 8B). This M1 stimulation led to whisking and nose twitches with shorter average latencies than oDAS (Figures 8B, 8C, S8A, and S8B). We also tested transient stimulation of wM1 neurons with a brief high frequency train of blue light pulses (5 ms, 4 times at 100 Hz), which resulted in transient whisker and nose movements resembling cue-locked orofacial movements (Figures 8D, 8E, S8C, and S8D). The average latencies of M1-driven transient whisker and nose movements were significantly shorter than those of cue-locked movements (Figures 8D and 8E). Thus, wM1-driven orofacial movements exhibited shorter onset latencies than orofacial movements after sound presentation or oDAS. Next, we explored whether wM1 stimulation could replicate certain aspects of orofacial movements. Whisker movements induced by sustained stimulation (1 s, 20 Hz, 20 times) exhibited greater vigor and duration compared to those induced by transient stimulation (100 Hz, 4 times), differentiable through a machine learning model based on protraction amplitude and ΣWM (Figure 8F). Our model successfully distinguished wM1-driven transient whisker movements from oDAS-induced motion (Figure 8G). However, the models were unable to differentiate between wM1-driven transient whisker movements and cue-locked whisker movements (Figure 8G). Through various stimulation protocols on wM1, we discovered a specific pattern involving 5-ms pulses at 50 Hz for 25 times that induced whisker movements akin to oDAS-induced but not cue-locked motion (Figure 8G). These results suggest that wM1 stimulation with different patterns can effectively elicit orofacial movements similar to those observed during the sound-oDAS pairing task.

**Figure 8.**
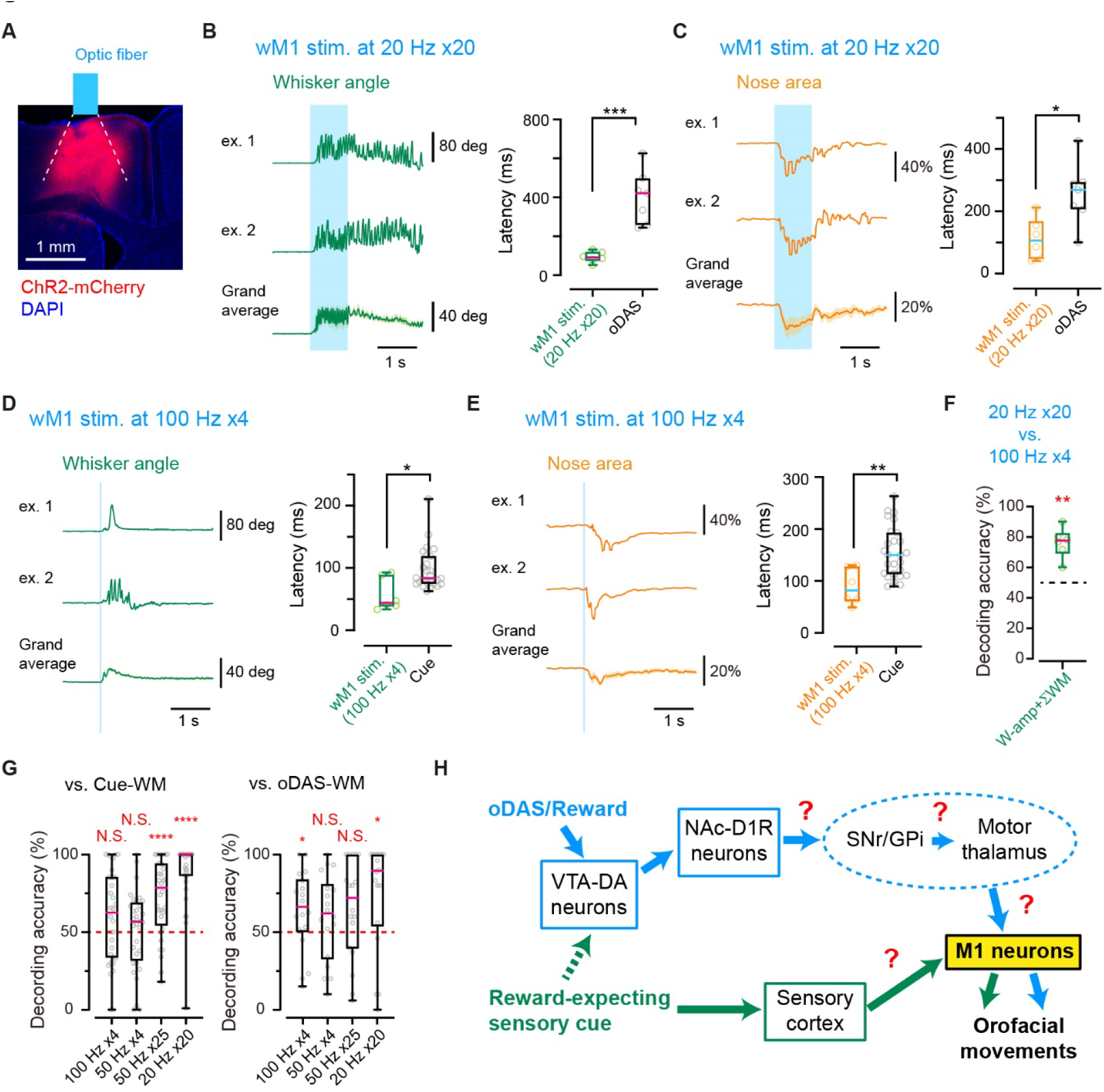
Optogenetic stimulation of wM1 induced whisker and nose movements. (A) An epifluorescence image of a coronal section indicating the AAV injection site at wM1. A schematic of the range of blue light illumination (dotted lines) with an optic fiber is superimposed. (B) Left, example and grand average traces of the whisker position upon photostimulation (5 ms, 20 pulses at 20 Hz) of wM1 expressing ChR2 (n = 6 mice). Right, Onset latency of the whisker movement upon wM1 photostimulation and oDAS (same data as Figure 2J, 20 Hz x20). (C) Same as (B) but for the nose area (left) and the onset of nose movement (right). (D) Same as (A) but with a transient wM1 photostimulation (5 ms, 4 pulses at 100 Hz; n = 6 mice). The onset latency of whisker movements is compared to that of the cue-locked movements (same data as Figure 3G, D2) (E) Same as (C) but with the transient stimulation. (F) Decoding accuracy by a model based on the whisker protraction amplitude (W-amp) and ΣWM to discriminate between wM1 stimulation with 20 pulses at 20 Hz and that with 4 pulses at 100 Hz. (G) Decoding accuracies by models based on W-amp and ΣWM for distinguishing wM1 driven whisker movements with different stimulus protocols and those at the cue onset of the sound-oDAS task on day 2 (left, Cue-WM) or during oDAS (right, oDAS-WM). (H) Schematic of presumable neural circuits for triggering uninstructed orofacial movements. SNr: substantia nigra pars reticulata, GPi: globus pallidus interna. Medians of the box plots are shown in cyan. Shadows of grand average traces: ± SEM. Open circles correspond to individual data.****p < 0.0001, **p < 0.01, *p < 0.05, N.S., not significant, unpaired *t* test in (B), (C), and (F), Mann-Whitney *U* test in (D), or one-sample *t* test vs. the chance level (50%) in (F) and (G), except for 50 Hz x25 and 20 Hz x20 vs. Cue-WM and 20 Hz x20 vs. oDAS-WM in (G) with one-sample Wilcoxon test. See also Figure S8.

## DISCUSSION

Through high-speed filming of orofacial movements of head-restrained mice performing reward-based learning tasks, we have characterized uninstructed facial movements at the timings of expectation and acquisition of a reward. Previous studies showing task-aligned uninstructed orofacial movements^5–8, 10–13^ used tasks involving motor actions for execution, such as holding handles, locomotion, and licking. The uninstructed orofacial motions during a task typically occur along with goal-directed actions. Using a Pavlovian learning paradigm, however, we demonstrated that uninstructed orofacial movements would become task aligned even without any need for motor action.

In our stimulus-reward association tasks, the cue-locked whisker protraction is induced immediately after reward-predicting cue presentation, almost coincident with the starting of anticipatory or goal-directed licking (Figure 1). Similar cue-locked whisker protraction can be elicited even without licking in our stimulus-oDAS pairing task (Figure 3). The mean amplitude of cue-locked whisker protraction increases as the mice learn the tasks. Such generality and scalability of the behavior suggest that the cue-locked whisker protraction may not merely be a reflexive movement upon sensory stimuli but rather influenced by the expectation of obtaining a reward^12, 14, 15^. A quick nose twitch is often coupled to the cue-locked whisker protraction in the stimulus-oDAS pairing task. These highly reproducible cue-locked orofacial movements appeared only transiently, followed by more random movements in the whisker and nose at the later phases of the cue presentation. Our observations are in line with the studies showing increased body and facial activities, including enhanced locomotion and sniffing at a higher frequency upon reward expectation^12, 43–46^. A classical study^47^ described a stereotypical movement sequence during a sniffing cycle, starting with head fixation, whisker protraction, a brief inhalation, and retraction of the tip of the nose. Therefore, the cue-locked whisker protraction and quick nose twitch described in the present study may correspond to a single sniffing cycle.

Facial movements evoked by oDAS were restricted mainly in the orofacial parts (Figure 2B). The oDAS-induced and cue-locked whisker movements have distinct properties in amplitude, vigor, and time course (Figures 3H and 3I): oDAS-induced motions have longer latency, are more persistent than the cue-locked whisker protraction, and generally have a protracted set point, similar to exploratory whisking^30–32^ as well as active whisking upon water reward except for licking-associated oscillatory movements (Figure 3J). Nose twitches also accompany the oDAS-induced whisker movements, suggesting the occurrence of sniffing^44, 47^. The movements of whiskers in response to oDAS display scalable positive relationships with both the stimulus frequency (Figure 2) and light intensity (Figures S2A– S2C), indicating a connection to reward-prediction error signals in the brain^19, 26–28, 48^. The significant kinetic differences in oDAS (Figure 2J)- and cue (Figure 3G)-induced motions suggest that different neuronal mechanisms drive these two types of orofacial movements.

Whisker movements are driven by facial muscles innervated by cholinergic motor neurons in the lateral facial nucleus. These whisker motor neurons receive synaptic inputs from premotor neurons, which are distributed in multiple areas within the brain stem, the midbrain, and the neocortex^49, 50^. Although there are significant kinetic differences in oDAS and cue-induced motion, whisking and nose movements often occur simultaneously. The respiratory centers in the ventral medulla that coordinate whisking and sniffing^17^ are thus likely involved in uninstructed orofacial movements. Our perturbational analyses indicate that VTA-DA neurons could directly or indirectly regulate the activity of these premotor circuits. Accumbal D1Rs are indispensable for the oDAS-induced orofacial movements (Figure 4). D1R-expressing GABAergic neurons in the NAc (NAc-D1R neurons) have outputs to the substantia nigra pars reticulata (SNr) which contains GABAergic neurons projecting to motor-related nuclei of the thalamus (motor thalamus). Activation of NAc-D1R neurons by oDAS thus leads to activation of the motor thalamus and, subsequently, the motor cortex. We found that neuronal activities in wM1 are essential for generating oDAS induced whisker movements (Figure 6) and identified specific neuronal populations that signal oDAS-induced whisker movements in wM1 (Figure 7). The wM1 strongly innervates brain stem reticular nuclei containing whisker premotor neurons^30, 31^, which may form a central pattern generator for rhythmic whisking^17, 51^. Optogenetic stimulation of wM1 could induce whisker and nose movements similar to oDAS but with a shorter latency (Figure 8). Taken together, our findings suggest that multi-nuclei transmission of signals from VTA to wM1 might facilitate the uninstructed orofacial movements in mice (Figure 8H). However, we cannot exclude the possibility that signals from NAc-D1R neurons through pathways independent of wM1 might also contribute to the activation of motor neurons in the facial nucleus.

The transient whisker protractions upon reward-predicting cues are reminiscent of transient firings of VTA DA neurons and subsequent DA release in the NAc immediately after reward-predicting cue presentation^26–28, 52^. DA released in the NAc would activate D1R expressing neurons generating the signal flow toward wM1, as discussed above. However, our results show that D1Rs do not mediate the cue-locked orofacial movements (Figure 5). The latency of cue-locked whisker protraction (∼100 ms) is much faster than oDAS-induced motion (∼400 ms). Therefore, learning a stimulus-reward association might recruit a cortical sensorimotor-coupling for reward expectation signaling, thereby activating wM1 earlier than the signals through NAc-D1R neurons. Transient stimulation of wM1 could evoke transient whisker movements with protraction, similar to but with a shorter latency than the cue-locked movements (Figure 8). The oDAS-induced and cue-locked movements are thus likely driven by different signal pathways dependent and independent on NAc-D1Rs, but these signals would converge into wM1 for facilitating orofacial movements (Figure 8H). Differences in the onset latencies (Figure 3G) and the sensitivity to a D2R antagonist (Figures 5E–5G) suggest that the neuronal mechanisms of cue-locked nose twitches might differ from those of the cue-locked whisker protraction. In fact, the magnitudes of these two movements were only weakly correlated (Figure S3C). Given that only minor fraction of wM1 neurons specifically represents nose movements (Figure S6E), uninstructed nose movements might be triggered by different M1 areas, including the subregion of orofacial M1 more closely related to nasal movements^33^, which could be inhibited by our pharmacological experiments (Figure 6).

Our findings provided insights into the neuronal mechanisms of uninstructed, task-aligned orofacial movements. We found that different neuronal populations in the wM1 area signal cue-locked, transient whisker protractions and oDAS-aligned, active whisking. This finding supports previous studies^32, 34, 36^ that indicate different cells within the wM1 region represent various aspects of whisker movements. Given the strong correlation between orofacial movements and brain-wide neuronal activity^4–7, 9^, it is likely that the activity of these wM1 neurons not only contributes to orofacial movements during reward-based behaviors but also has a broader impact on overall brain processing.

Our study might naturally prompt an intriguing question: why do mice move their whiskers and nose at the moments of reward anticipation and acquisition? We speculate that when mice feel rewarded, they could be attempting to gather sensory information from their immediate surroundings. This action could be a mechanism for memorizing the environment, leading to rewarding experiences. Alternatively, these orofacial movements may serve as perceptible signals to conspecifics conveyed through social facial touches. The transmission of these reward-related internal states between individuals could play a vital role in shaping the prosocial behaviors of animals. Therefore, it would be of critical importance to further investigate the role of these uninstructed orofacial movements in social communication.

## METHODS

### Animals

All experimental procedures were approved by the Swiss Federal Veterinary Office (for whisker detection task experiments), the institutional review board of the Research Institute of Environmental Medicine, Nagoya University, and the Institutional Animal Care and Use Committee of Fujita Health University. For optogenetic experiments, we used DAT-ChR2 mice by crossing DAT-IRES-Cre mice [B6.SJL-*Slc6a3^tm1.1(cre)Bkmn^*/J] with Ai32 mice [B6;129S-*Gt(ROSA)26Sor^tm32(CAG–COP4∗H134R/EYFP)Hze^*/J]^54^, Ai32 mice as control or DAT-IRES Cre mice with AAV injected (see below). For behavioral experiments using water reward based learning tasks (Figure 1), we used adult male C57BL/6J wild-type mice. Mice were at least 6-week-old at the time of head-post implantation (see below). Mice were kept in a shifted light/dark cycle (light 0 p.m. to 0 a.m. in Nagoya University and Fujita Health University; light 7 p.m. to 7 a.m. in EPFL) in ventilated cages at a temperature of 23 ± 3°C with food available ad libitum. Behavioral experiments were performed during the dark period. Water was restricted during the training of water reward-based learning tasks. All mice were weighed and inspected daily during behavioral training and given wet food, so that mouse weight was kept to be more than 80% of the initial value.

### Plasmids

For AAV production, pAAV-Ef1a-DIO-bReaChES-TS-eYFP was obtained from K. Deisseroth (Stanford University). For CRSPR-Cas9-mediated gene editing, we used a plasmid containing a saCas9 encoding sequence with *Drd1a*-targeting short guide RNA^42^ (target sequence: 5’-GTATTCCCTAAGAGAGTGGA-3’)^41^: pairs of *Drd1a*-targeting oligo DNAs (forward, 5’-CACCGGGTATTCCCTAAGAGAGTGGA-3’, reverse, 5’-AAACTCCACTCTCTTAGGGAATACCC-3’) were ligated into a plasmid pX601-AAV-CMV::NLS-SaCas9-NLS-3xHA-bGHpA;U6::BsaI-sgRNA (pX601, Addgene #61591) digested with BsaI (R0535, New England BioLabs). We constructed a plasmid with a negative-control saCas9 guide RNA by shuffling bases of the *Drd1a*-targeting guide RNA sequence (target sequence: 5’-TCAATAATGAGGTGGTCCGA-3’): the pX601 was linearized with BsaI, and a pair of oligonucleotides (Forward, 5’-CACCGTCAATAATGAGGTGGTCCGA-3’; Reverse, 5’-AAACTCGGACCACCTCATTATTGAC-3’) was ligated with T4 DNA Ligase (M0202, New England BioLabs).

### Viral production

To produce AAV vectors for axonal stimulation experiments (Figures 4E and 4F), HEK293 cells were transfected with vector plasmids including pAAV encoding bReaChES or hrGFP together with pHelper, and pAAV-RC (serotype 9 or DJ), using a standard calcium phosphate method. After three days, transfected cells were collected and suspended in lysis buffer (150 mM NaCl, 20 mM Tris pH 8.0). After four freeze-thaw cycles, the cell lysate was treated with 250 U/ml benzonase nuclease (Merck) at 37 °C for 10–15 min with adding 1 mM MgCl_2_ and then centrifuged at 4 °C at 1753×g for 20 min. AAV was then purified from the supernatant by iodixanol gradient ultracentrifugation. The purified AAV solution was concentrated in PBS via filtration and stored at −80°C.

For *in vivo* genome editing experiment (Figures 4C and 4D), HEK293 cells were transfected with vector plasmids including pAAV encoding *Drd1a*-targeting guide RNA or control guide RNA, together with pHelper, and pAAV-RC (serotype 9), using a standard calcium phosphate method. Transfected cells were collected after three days and suspended in Hank’s Balanced Salt Solution (Merck). The AAV-containing cell lysate was treated with 250 U/ml benzonase at 37 °C for 30 min without adding MgCl2, and then centrifuged three times at 17,800×g for 10 min at 4°C with the supernatant after each centrifugation used for the next centrifugation. The final supernatant was then aliquoted and stored at −80°C.

### Heteroduplex cleavage assay

To examine the off-target effect of our saCas9-*Drd1a*-gRNA, we performed T7 endonuclease I (T7EI) assay using Alt-R Genome Editing Detection Kit (Integrated DNA Technologies). The saCas9-*Drd1a*-gRNA plasmid or a control hrGFP-expressing plasmid (pAAV-hrGFP Vector, #240074, Agilent) was transfected into NIH/3T3 cells in a 24-well plate using jetOPTIMUS (PolyPlus). After 72 h post-transfection, genomic DNA was isolated using Guide-it Mutation Detection Kit (Takara). DNA fragments flanking the targeted *Drd1a* locus and off-targets (Figure S4C) were then amplified from the purified genomic DNA with PCR. Positive and negative control DNA fragments provided by AltR-Genome Editing Kit were also amplified with PCR. PCR products were denatured and hybridized, digested at 37°C for 60 min with T7EI, and analyzed by electrophoresis in 2% agarose gel stained with ethidium bromide. The gel images were obtained with a transilluminator (E-BOX VX2, Vilber Lourmat).

### Animal preparation and surgery

Mice were anesthetized with isoflurane (3.0–3.5% for induction, 1.0–1.5% for maintenance) and head-fixed on a stereotactic device using ear bars or a nose clamp. Body temperature was maintained at ∼37°C by a controlled heating pad. An ocular ointment was applied over the eyes to prevent drying. The scalp was cut open to expose the skull. For *in vivo* genome editing (Figures 4C and 4D), AAV harboring *Drd1a*-targeting guide RNA or control guide RNA was bilaterally injected into four sites targeting the NAc (AP: 1 mm, ML: ±1.2 mm, from Bregma, depth: 3.8 mm; and AP: 1.5 mm, ML: ±1.2 mm, from Bregma, depth: 4.0 mm) through small craniotomies (diameter, <∼0.5 mm). The injection volume was 500 nl per site. For the experiment with optogenetic axonal stimulation (Figures 4E and 4F), AAV-Ef1a-DIO-bReaChES-TS-eYFP (titer: 6.0 × 10^13^ copies/ml) or AAV-CMV-DIO-hrGFP (titer: 6.0 × 10^12^ copies/ml) was injected bilaterally into the VTA (AP: −3.0 mm, ML: ±0.5 mm, from Bregma, depth: 4.2 mm) of DAT-IRES-Cre mice. The injection volume was 200 nl per site. For the experiment with optogenetic stimulation of the SNc-DA neurons (Figures S2E–S2G), AAV9-Ef1a-DIO-hChR2(E123T/T159C)-EYFP (Addgene #35509, obtained from Addgene; the original titer: 2.2 × 10^13^ copies/ml, diluted to 1/10) or AAV9-Ef1a-DIO-EYFP (Addgene #27056, obtained from Addgene; the original titer: 2.7 × 10^13^ copies/ml, diluted to 1/10) was injected into the left SNc (AP: -3.1 mm, ML: 1.2 mm, from Bregma, depth: 4.1 mm) of DAT-IRES-Cre mice. The injection volume was 200 nl. For optogenetic stimulation of wM1, AAV9-CaMKIIa-hChR(E123T/T159C)-mCherry (Addgene #35512, obtained from Addgene; the original titer, 3.0 × 10^13^ copies/ml, diluted to 1/10) or AAV9-Ef1a-mCherry (Addgene # 114470, obtained from Addgene; the original titer, 1.0 × 10^13^ copies/ml, diluted to 1/10) was injected unilaterally into the left wM1 (AP: 1.0 mm, ML: 1.0 mm, from Bregma, depth: 0.85 mm and 0.35 mm) of wild-type C57BL6/J mice. The injection volume was 200 nl per site. With the scalp sutured, the mice were returned to their home cages for at least three weeks after the AAV injection before behavioral experiments.

Prior to behavioral experiments, mice were implanted with a light-weight metal head holder^37^. For optogenetic stimulation, pharmacological perturbation, or silicone probe recordings in left wM1, a chamber was made on the left hemisphere by building a wall with dental cement, and a thin layer of glue was applied over the exposed skull. For optogenetic stimulation of deep brain regions, we used an optical fiber (0.4 mm diameter) attached to a stainless steel ferrule (CFM14L05, Thorlabs) which was implanted over the left NAc (AP: 1.25 mm, ML: 1.5 mm, from Bregma, depth: 3.8 mm) or the left VTA (AP: −3.0 or −3.3 mm, ML: ±0.3 or 0.5 mm, from Bregma, depth: 4.0 mm). The optical fiber cannula was permanently cemented to the skull. For optogenetic stimulation of wM1, we made a small craniotomy over the left wM1 and placed the optical fiber over the craniotomy.

### Behavioral tasks

#### Whisker detection task

To examine the orofacial movements of mice performing a whisker detection task^10^, we analyzed the data obtained from the mice used for recording in our previous paper^37^. The procedure of the whisker detection task was previously described^37^. All whiskers except for the right C2 whisker were trimmed before experiments. We applied a brief (1 ms) magnetic pulse to elicit a vertical deflection of the right C2 whisker transmitted by small metal particles glued on the whisker. The mice under head-fixation were trained to learn the availability of water reward within a 1-s window after the whisker stimulation. The whisker was filmed at 500 Hz with a high-speed camera. The behavioral signals from the lick sensor together with TTL signals to control the water valve and the electromagnetic coil were recorded through an NI board. In some recordings, it was impossible to extract the whisker position due to the metal particles covering the basal part of the whisker. Therefore, we only included the data where the basal part of the whisker was partially exposed. Novice mice had no prior task experience and showed a low hit rate (31.1 ± 4.2%, n = 6 mice). Expert mice analyzed in this study were trained for the task in 8–17 daily sessions before the recording day and exhibited a high hit rate (79.2 ± 3.6%, n = 6 mice).

#### Auditory Go/No-Go task

At least three days after implantation of the metal head holder, adult male C57Bl6/J mice started to be water-restricted. The mice were adapted to head restraint on the experimental setup through initial training to freely lick the water spout for receiving a water reward (3–5 sessions, one session per day). The mice were then taught, through daily training sessions, to associate a pure tone (2 s) with water availability within a 1-s window after the offset of tone presentation. Trials were started with a 3 kHz tone (“Go” cue) or a 15 kHz tone (“No Go” cue) following random inter-trial intervals ranging from 3 to 9 s. In some experiments, the “Go” tone and “No-Go” tone were swapped. If the mice licked in the 2 s preceding the time when the trial was supposed to occur, then the trial was aborted, and a subsequent trial started. Lick was detected with a piezo sensor attached to the water spout. After each training session, 1.0–1.5 g of wet food pellet was given to the mouse to keep its body weight above 80% of the initial value. Behavioral control was carried out using a custom-written program on Python interfaced through Arduino Uno. All whiskers except for the right C2 whisker were trimmed before experiments. The whisker was filmed at 200 Hz with a high speed camera. Behavioral data were acquired using an NI board. Novice mice were defined by poor performance (daily d’ < 1.0). These mice were typically on the first day of task training and showed an average hit rate of 33.7 ± 5.5% (n = 9 mice). Expert mice were defined as those keeping a high performance (daily d’> 2.5) for five successive days. These mice showed an average hit rate of 94.1 ± 1.4% (n = 7 mice). One mouse underwent a reversal-learning: after the mouse learned the task with a 3-kHz tone as a Go-cue, then we swapped the Go and No-Go tones. The mouse successfully learned the swapped association, and the data were pooled into the dataset as a different experiment from the initial learning.

### Optogenetics

#### Optogenetic stimulation in head-fixed mice

All the behavioral experiments using optogenetic stimulation followed the habituation of mice to head-fixation for three days. The duration of habituation was ∼15 min on the first day, ∼30 min on the second day, and ∼60 min on the third day. All whiskers were trimmed except the left C2 whiskers or the left and right C2 whiskers before the experiments. For optogenetic stimulation of DA neurons, blue LED (470 nm; M470F3, Thorlabs) light was applied over the VTA or SNc of DAT-ChR2 mice or control Ai32 mice through the implanted optic fiber. For stimulating bReaChES/hrGFP-expressing DA axons in the NAc, green LED (530 nm; M530F2, Thorlabs) light was applied over the NAc through the implanted optic fiber. For stimulating ChR2-expressng neurons in wM1, the blue LED light was applied through the craniotomy made over the left M1. 20 pulses of 5-ms light stimuli were applied at 20 Hz unless otherwise noted. The light power at the fiber tip was 15.8 mW for the blue light and 6.6 mW for the green light unless otherwise noted. The frontal mouse face, including its nose and whisker(s), was filmed at 500 Hz from above with a high-speed camera. Through immunostaining (see below), we routinely checked the location of implanted optical fiber and discarded data when the fiber tip was found to be more than 0.5 mm away from the target or inserted too deep damaging the target brain structure.

#### Sound-oDAS pairing conditioning

For the Pavlovian sound-oDAS conditioning, we used the mice that experienced oDAS. We habituated the mice to a 15-kHz tone presentation (5 s) by being exposed to the tone stimuli 20 times with random intervals of 60–120 s under head fixation on the day before starting the conditioning. For the conditioning, we presented to the mice the 15-kHz sound cue (5 s) paired with oDAS (5 ms, 20 times at 20 Hz) at the last 1 s of the cue. The inter-trial interval was set randomly for each trial to be from 180–240 s. A custom-written program on Python interfaced through Arduino Uno controlled the timings of sound cue presentation and light stimulation, which were synchronized with filming the mouse’s frontal face from above. The mice experienced ∼20 paired stimuli in each daily session.

### Pharmacological perturbation

To examine the effect of a D1R antagonist on orofacial movements, we first calculated the injection volume of SCH23390 (0.3 mg/kg) or raclopride (3 mg/kg) for each experimental mouse and injected saline of the calculated volume intraperitoneally into the mouse. The behavioral recording was started 15 min after injection. After the recording session with saline injection, the mice were subjected to an IP injection of the D1R/D2R antagonists, and the behavioral recording was resumed 15 min after injection.

To examine the effect of wM1 inactivation on orofacial experiments, we first made a small craniotomy over wM1 of DAT-ChR2 mice under anesthesia. After recovery from anesthesia, the mouse was head-fixed to the experimental setup, and the behavior was recorded. Subsequently, 100 nl of Ringer’s solution with or without muscimol (5 mM) was injected into at 900, 700, 500, 300, and 100 µm each below the surface using a glass pipette inserted through the craniotomy. A small volume of Chicago Sky Blue was mixed in the muscimol solution. The whole injection period was 20–30 min. The behavioral tests were resumed after the injection pipette was slowly withdrawn.

### Immunostaining

After behavioral and electrophysiological experiments, we routinely performed transcardial perfusion of the mice with 4% paraformaldehyde (PFA). The brain was removed and incubated with 4% PFA solution overnight for post-fixation. The fixed brain was then kept in phosphate buffer (PB) until further processing. The fixed brain was sectioned into coronal slices on a vibratome (section thickness: 100 μm). For immunostaining of brain slices, the slices were washed three times with a blocking buffer containing 1% bovine serum albumin and 0.25% Triton-X in phosphate buffer saline (PBS) and then incubated with primary antibodies (anti-tyrosine hydroxylase, rabbit polyclonal, 1:1000, Merck Millipore; anti-D1R, guinea pig polyclonal, 1:200, Frontier Institute; anti-GFP, mouse monoclonal, 1:1000, Fujifilm Wako Chemicals) in the blocking buffer overnight at 4 °C. The slices were then washed three times with the blocking buffer and then incubated with secondary antibodies (CF594-conjugated donkey anti-rabbit IgG, 1:1000, Biotium; CF488A-conjugated goat anti mouse IgG, 1:1000, Biotium; CF594-conjugated donkey anti-guinea pig IgG, 1:1000, Biotium) in the blocking buffer for 1–2 h at room temperature (RT). Cellular nuclei were stained by incubation for 10–15 min with DAPI (2 μM in PB) at RT. The stained samples were mounted using DABCO and observed under a fluorescence microscope (BZ-9000, Keyence). Images were saved using BZ-X Viewer (Keyence).

### Electrophysiology

A small craniotomy was made over the left wM1 (1 mm anterior, 1 mm lateral from Bregma). Extracellular spikes were recorded in head-fixed mice using a silicon probe (A1x32-Poly2-10mm50s-177, NeuroNexus) with 32 recording sites along a single shank, covering 775 mm of the cortical depth. The probe was lowered gradually until the tip was positioned at a depth of 1.0–1.1 mm under the wM1 pial surface. The craniotomy site was then covered with 1.5% agarose dissolved in Ringer’s solution. Neural data were filtered between 0.5 Hz and 7.5 kHz, amplified using a digital head-stage (RHD2132, Intan Technologies), and digitized with a sampling frequency of 30 kHz. The digitized neural signal was transferred to an acquisition board (Open Ephys) and stored on an internal HDD of the host PC for offline analysis. The TTL pulses for the sound and light stimulation were also recorded through the Open Ephys acquisition board. Spiking activity on each probe was detected and sorted into different clusters using Kilosort (https://github.com/cortex-lab/KiloSort). After an automated clustering step, clusters were manually sorted and refined using Phy (https://github.com/cortex-lab/phy). Only well-isolated single units (174 units) were included in the dataset.

## Quantification and statistical analysis

### Motion energy analysis

Mouse facial movements were quantified by analyzing the side-view (Figure 2B) and top view (Figure 2C) movies. Motion energy was computed as the absolute value of the difference of consecutive frames obtained during oDAS (for 1 s) and at the baseline (0–1 s before the stimulus onset). The motion energy heat-map is shown as the difference between motion energy during oDAS and at the baseline in each pixel. For quantification (Figure 2B), we set polygonal regions of interest around the ear or orofacial area and calculated the mean change in motion energy evoked by the oDAS by comparing values during oDAS and at baseline.

### Analysis of whisker movements

Movements of the right C2 whisker were quantified offline with ImageJ using an open source macro (https://github.com/tarokiritani/WhiskerTracking) and the data were analyzed with Igor Pro or MATLAB. The total whisker movements (ΣWM) were calculated as the cumulative whisker movements during oDAS (Figures. 2, 4, 6, S2, S3D, S6, and S8), at 0– 1 s after the reward time window for the data in the reward-based learning tasks (Figure 1) or at 0–0.5 s after the cue onset in the sound-oDAS pairing task (Figure 5, S3B, S5B, and S6A). Data with the prominent whisking starting during the 1-s period before the onset of oDAS (Figures 2, 4, 6A–6D, S2, S4, S5A, S5C, and 8) or the conditioning stimulus (Figures 3, 5, 6E–6H, 7, S3A–S3C, S5B, S6, and S7) were excluded from the analysis. When the ΣWM during oDAS exceeded 90 deg, the data were defined as epochs with whisking induced (WM+). A cue-locked whisker protraction was defined as a whisker protraction of more than 5 deg above baseline within 0.5 s after the onset of cue presentation. The onset time of whisker movement or whisker protraction was taken as the time at which the whisker angle exceeded 1 deg above baseline, among these epochs with whisker movement or cue locked whisker protraction.

### Analysis of nose movements

We quantified nose movement using the markerless video tracking software, DeepLabCut^38^. As illustrated in Figure 2G, we manually annotated the five points of the top-view nose (left and right anterior/posterior edges and nose tip) using 60–340 frames per movie to train a deep neural network. An estimated position with a low likelihood (<0.8) was omitted and replaced with a pixel value obtained using linear interpolation of neighboring values^55^. The nose area was the area of the pentagon formed by the estimated five points. The nose length in the anterior-posterior axis was taken as the distance between the nose tip and the middle point of the posterior nose edges. The lateral movement of the nose tip was the displacement of the horizontal coordinate of the nose tip quantified as the percentage with the average anterior-posterior nose length taken as 100%. The onset of lateral nose tip movement was defined as the time point where the trace exceeded three times of standard deviation of the baseline (0–1 s before the stimulus/cue onset). The latency of nose movement was defined as the time difference between the stimulus/cue onset and the onset of lateral nose tip movement. The data with whisking at the pre-stimulus period (1 s) were excluded from the analysis of nose movement.

### Machine learning models

We utilized machine learning models to decode trial categories (“Hit” or “CR”) of the auditory Go/No-Go task using whisker time plots. We selected “Hit” and “CR” trials from the dataset while excluding “Miss” and “False Alarm (FA)” trials. We filtered the data for each trial with a band-pass filter (4–20 Hz) and transformed it into a power spectrogram using short-term Fourier transformation with 200-ms or 100-ms bins. The power spectrogram was then extracted at 4-20 Hz and linearly normalized so that the sum of powers equaled one in each time bin. We created power spectrum matrices with 240 dimensions (24 frequency domains x 10 time domains) for the data during the cue period (2 s) or 0-1 s after the reward time window (after RA). Among the data from seven expert mice, we pooled the data from six mice as training data, excluding the one from one mouse as test data, and applied the principal component analysis (PCA) on MATLAB to make a PC space with two dimensions. We used this PC space to obtain the first and second PC scores for each trial of the test data and plotted them (Figure S1C). We constructed a logistic regression model using the PC plot obtained from training data, which outputs a likelihood (minimum: 0, maximum: 1) of being “Hit” for each trial of the test data. The trials with a likelihood of more than 0.5 were predicted as “Hit” trials, and otherwise, the trials were predicted as “CR” trials. The decoding accuracy is a fraction of the number of correct predictions over the total trial number of the test data. We repeated the same procedure for the other six mice (a process called leave one-subject-out cross-validation).

We also used leave-one-subject-out cross-validation to train logistic regression models for discriminating between cue-locked and reward-aligned whisker movements during the auditory Go/No-Go task (Figure 1H), oDAS-aligned and cue-locked orofacial movements during the sound-oDAS pairing task (Figure 3H and 3I), whisker movements during the sound-oDAS pairing task and those during the auditory Go/No-Go task (Figure 3J), whisker movements with different wM1 stimulus patterns (Figure 8F), and wM1-driven whisker movements and those in the sound-oDAS pairing task (Figure 8G). As decoding features, we used the amplitude, total movements, and first PC score of multidimensional power spectrum matrices (see above) of whisker (W-amp, ΣWM, and W-spectral PC) and/or nose (N-amp, ΣNM, and N-spectral PC) obtained from 1-second traces during the cue onset, after RA, during oDAS or upon wM1 stimulation.

### Classification of single units

Single units recorded with the silicone probe were classified as fast-spiking (FS) putative interneurons or regular-spiking (RS) putative pyramidal cells based on their trough-to-peak time of average spike waveform. Single units with a trough-to-peak time < 0.35 ms were classified as FS cells, and units with a trough-to-peak time > 0.35 ms were classified as RS cells.

### Analysis of the correlation between spike rates and whisker behavior

We classified wM1 cells based on their correlation with whisker behavior during the sound oDAS pairing tests. We first categorized the trials into four categories (Figure 7A): “CL+OA trials” are trials with both cue-locked whisker protraction and oDAS-aligned whisker movements; “CL trials” are trials with cue-locked whisker protraction but without oDAS-aligned whisker movements; “OA trials” are trials with oDAS-aligned whisker movements but without cue-locked whisker protraction; and “No reward-related WMs trials” are trials with neither cue-locked whisker protraction nor oDAS-aligned whisker movements. We used different trial categories: “Cue-WP+ trials” are CL+OA and CL trials; “Cue-WP-trials” are OA and No reward-related WMs trials; “oDAS-WM+ trials” are CL+OA and OA trials; and “oDAS-WM-trials” are CL and No reward-related WMs trials. We defined the cellular categories as follows: “CL+OA cells” showed a significantly higher firing rate at the cue onset (1 s) in Cue-WP+ trials than in Cue-WP-trials, and also showed a significantly higher firing rate during oDAS in oDAS-WM+ trials than in oDAS-WM-trials; “CL cells” showed a significantly higher firing rate at the cue onset (1 s) in Cue-WP+ trials than in Cue-WP-trials, and exhibit similar firing during oDAS in oDAS-WM+ and oDAS-WM-trials; “OA cells” showed a significantly higher firing rate during oDAS in oDAS-WM+ trials than in oDAS-WM-trials, and exhibit similar firing at the cue onset (1 s) in Cue-WP+ and Cue-WP-trials. In Figure 7E, we plotted “CL correlation” as the Pearson correlation coefficients between spike rates and the maximal WP amplitudes during 1 s at the cue onset and “OA correlation” as the Pearson correlation coefficients between spike rates and the variances of the whisker position during oDAS. For motion-triggered PSTHs upon oDAS (Figure 7I), the data with whisking at 0–0.2 s before oDAS were excluded from the analysis.

### Statistics

All values are expressed as mean ± SEM except for the box plots in the figures. Statistical tests were performed using GraphPad Prism. The normality of data distribution was routinely tested. Analyses of two sample comparisons were performed using unpaired or paired *t* tests when each sample was normally distributed, or Wilcoxon signed rank test (paired) or Mann-Whitney *U* test (unpaired) when at least one of the samples in every two-sample comparison was not normally distributed. Tests for two-sample comparison were two-sided. Statistical analyses for multiple comparisons were carried out using one- or two-way ANOVA followed by Bonferroni’s multiple comparison tests or Dunnett’s multiple comparison tests vs. the control, unless otherwise noted. For correlation analysis, Pearson’s correlation coefficient (*r*) for normally distributed sets of data or Spearman’s rank correlation coefficient (rho) for randomly distributed data was calculated.

## Acknowledgements

We thank Dr. H. Kasai for providing DAT-IRES-Cre mice; Dr. K. Ono for providing Ai32 mice; Dr. K. Deisseroth for providing bReaChES plasmid; Drs. T. Hikida, Y. Ohmura and J. Dijkstra for discussion; J. Ozaki for collaboration at the early phase of the study; N. Fukatsu for helping with nose analysis; E. Charrière, S. Tsukamoto, A. Kambara, M. Jin, E. Imoto for technical assistance. This work was supported by JSPS KAKENHI grants (17H05744, 18K19496, 21H00215, and 23H04685) to TY; The Naito Foundation to TY; Takeda Science Foundation to TY; JNNS30 Commemorative Research Grant to JY.

## Author contributions

WL, TN, KM, JY, and TY designed the study. WL, KM, MK, TM, YM, and TY performed experiments and analyzed data. TN and JY developed machine learning algorithms and analyzed data. TD developed Matlab codes for motion energy analysis. AY provided plasmids and AAVs, and contributed to interpreting the results. HI and HA provided plasmids for in vivo genome editing, performed pilot experiments, and contributed to interpreting the results. TY and CCHP contributed to data collection related to the whisker detection task. WL and TY drafted and wrote the manuscript with inputs from other co-authors.

## Inclusion and diversity

We support inclusive, diverse, and equitable conduct of research.

---

**Figure S1.**
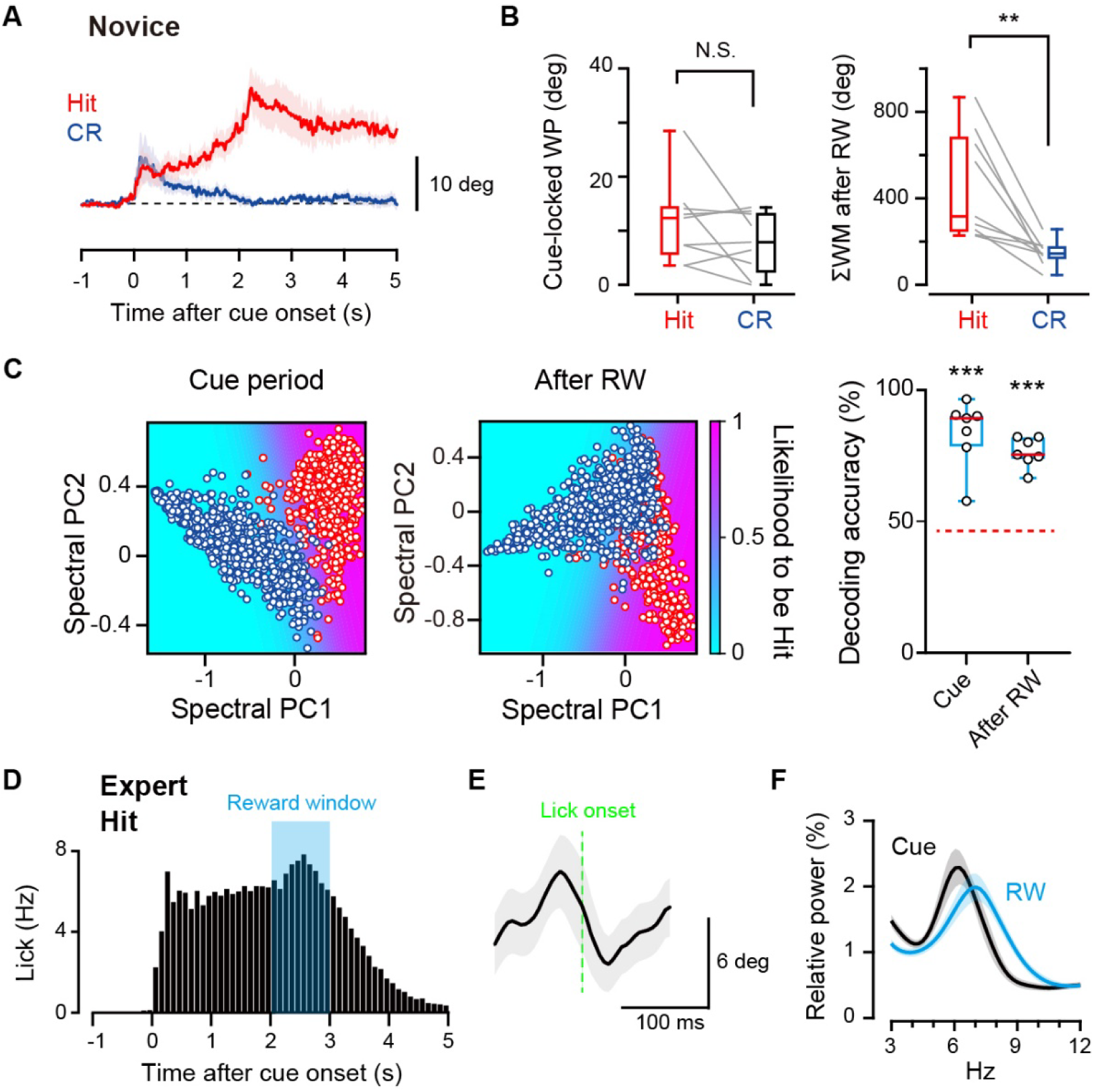
Additional analyses of orofacial behaviors during the auditory Go/No-Go task. (A) Grand averages of whisker movements in hit (red) and CR (blue) trials from 9 novice mice. (B) Data for each mouse (thin lines) and box plots for the amplitudes of the cue-locked whisker protraction (WP) (left) and total whisker movement (ΣWM) 0-1s after reward window (right), in hit (H) and miss (M) trials during the auditory Go/No-Go task, obtained from novice mice. **p < 0.01, N.S., not significant, paired *t* test. (C) Example principal component (PC) analyses of whisker time plots from an expert mouse during the cue period (left) and 0–1 s after reward time window (after RW, middle) of hit (red) and CR (blue) trials (see Methods). Decoding accuracies by the models (right) were significantly higher than the chance level for both cue period and after RW (n = 7 mice). The solid red line in the box plots indicates the median. The dashed red line indicates the mean chance level. ***p < 0.001, Dunnett’s multiple comparison test vs. the chance level. (D) Average lick PSTH during the auditory Go/No-Go task obtained from 7 expert mice. Light blue shadow: reward time window. (E) Lick-triggered average whisker trace at the cue period obtained from 7 expert mice. (F) Relative power spectra of whisker movements during the cue period (Cue) and the reward time window (RW) obtained from 7 expert mice. Thick lines and shadows in (A) and (E) indicate mean ± SEM. Thin lines in (B) and open circles in (C) indicate individual data.

**Figure S2.**
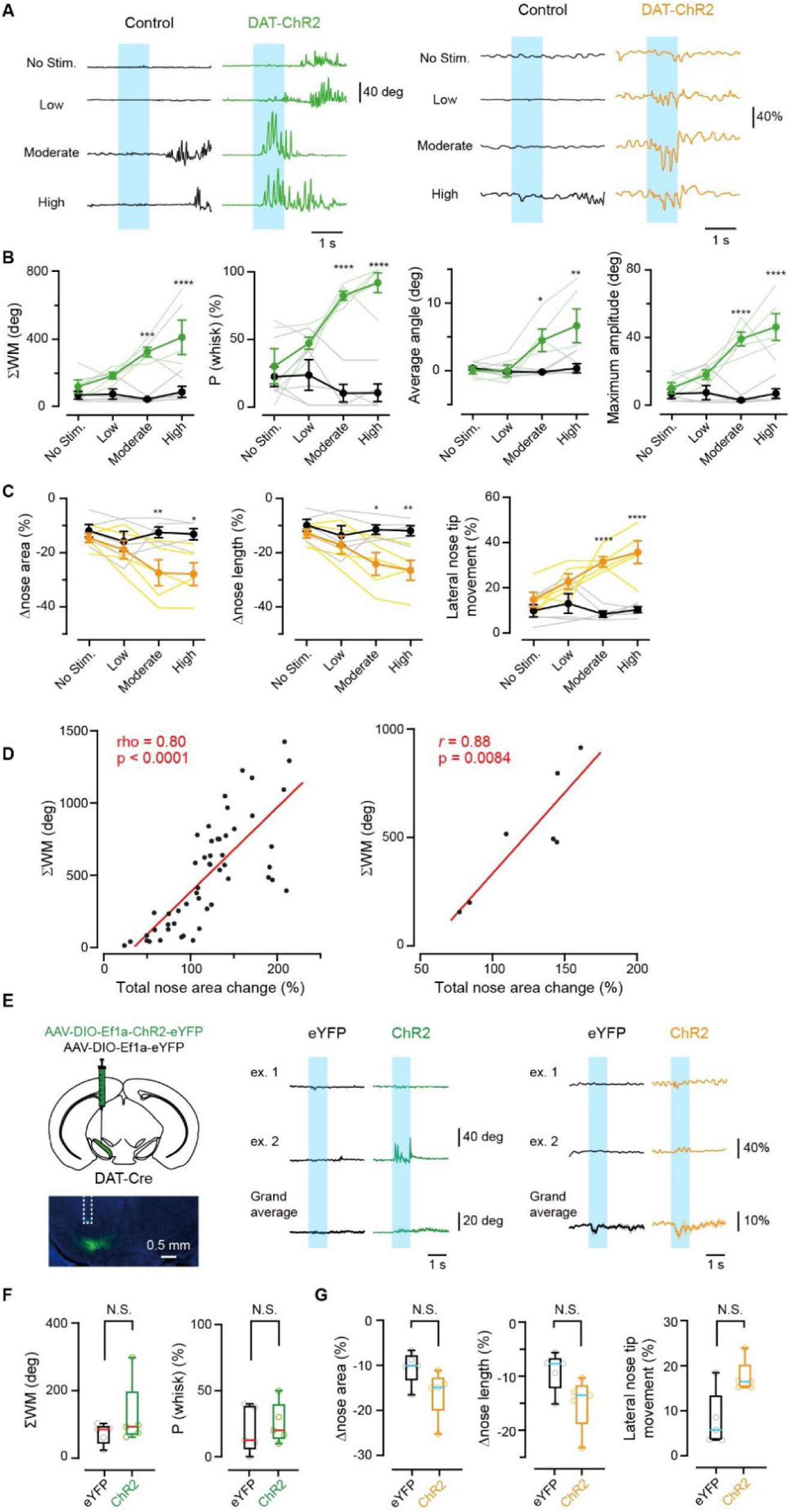
Additional analyses on oDAS-Induced orofacial movements. (A) Example traces of the whisker position (left) and nose area (right) upon oDAS (light blue shadow) with different photo-stimulus intensities, obtained from a control (black) and DAT-ChR2 (green) mouse. The light intensity at the fiber tip was 3.0 mW (low), 7.6 mW (moderate), or 15.8 mW (high). (B) Quantifications for whisker movements during oDAS with different stimulus intensities (n = 5 for each). (C) Same as (B) but for nose movements. (D) ΣWM as a function of the total change in the nose area for individual oDAS (5 ms x20, 20 Hz) trials (n = 52 trials, from 7 mice, left) or individual mice (n = 7 mice, right). We selected trials without prominent whisking starting 0-1 s before the stimulus onset. Red line: linear regression. (E) Left, schematic for the AAV injection into the SNc (top), and an epifluorescence image of a coronal section containing SNc and the trace of the inserted optical fiber (dashed line) (bottom). Green: ChR2-eYFP, blue: DAPI. Middle and right, example and grand average traces of the whisker position (middle) and the top-view nose area (right) obtained from ChR2- and eYFP-expressing mice (n = 5 for each group). (F) ΣWM and P(whisk) upon optogenetic stimulation of SNc-DA neurons in ChR2- and eYFP-expressing mice. (G) Same as (F) but for the parameters of nose movement. Medians of the box plots are shown in magenta or cyan. Lightly colored lines in (B) and (C) correspond to individual data. Shadows of grand average traces: ± SEM. Filled circles and error bars show mean ± SEM. \*\*\*\**p* < 0.0001, N.S.: not significant, Bonferroni’s multiple comparison test (Control vs. DAT-ChR2) in (B) and (C), unpaired *t* test (P(whisk) in (F) and Δnose area and Δnose length in (G)) or Mann-Whitney *U* test (ΣWM in (F) and lateral nose tip movement in (G)).

**Figure S3.**
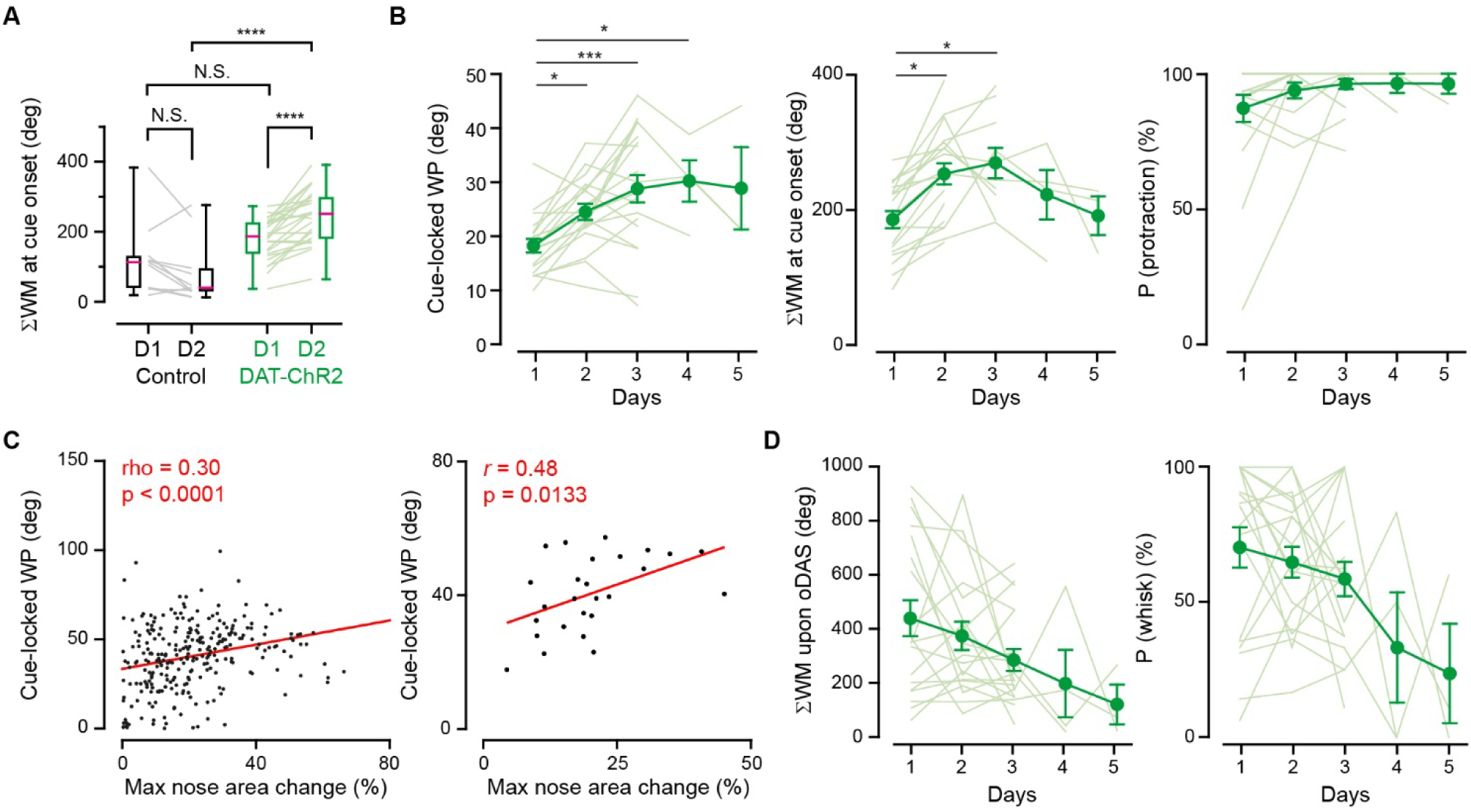
Additional analyses of whisker and nose movements during the stimulus oDAS association task. (A) ΣWM at the cue onset (0-0.5 s) at day (D)1 and 2 in control (black) and DAT-ChR2 (green) mice (control: *n* = 11 mice; DAT-ChR2: *n* = 26 mice). (B) Protraction amplitude (left), total amount (middle), and probability (right) of cue-locked whisker movement of DAT-ChR2 mice on day 1–5 during the sound-oDAS pairing conditioning (day 1–3, n = 19 mice; day 4, n = 4 mice; day 5, n = 3 mice). We performed pharmacological experiments on day 3–5 and data with saline injection (Figure 5) or before muscimol/Ringer injection (Figure 6) were plotted. We did not include data during wM1 recordings (Figure 7) nor the data from the mice used only for 2 days without pharmacological experiments. (C) The amplitude of cue-locked whisker protraction (WP) as a function of that of cue-locked nose area change for individual trials (left, n = 272 trials, from 26 mice, left) or individual mice (n = 26 mice, right) at day 2. We selected trials without prominent whisking starting 0-1 s before the stimulus onset. Red line: linear regression. (D) ΣWM (left) and the probability of whisking (right) of the same mice in (B) during oDAS in the sound-oDAS pairing conditioning. Filled circles and error bars show mean ± SEM. Lightly colored lines correspond to individual data. ****p < 0.0001, ***p < 0.001, *p < 0.05, N.S., not significant, Bonferroni’s multiple comparison test in (A) or Kruskal-Wallis test (vs. D1) in (B) and (D).

**Figure S4.**
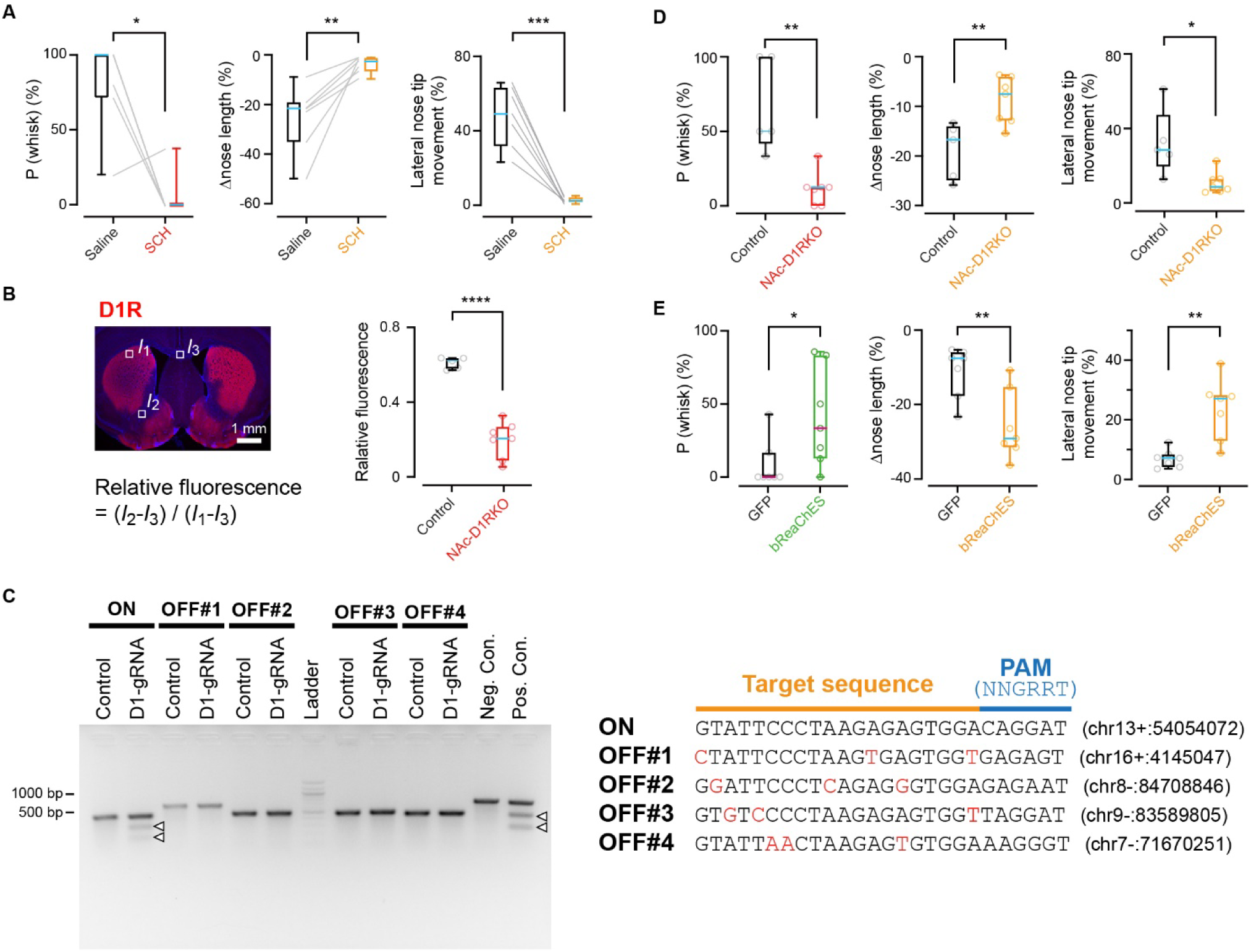
Additional analyses of perturbational experiments for oDAS-induced orofacial movements. (A) Left, data for each mouse (thin lines) and box plots for the parameters of whisker or nose movements in trials after IP injection of SCH23390 (red or orange) and saline (black) during oDAS (n = 7 mice). (B) Left, schematic for the fluorescence analysis. Average fluorescent signal intensity (FI) at the region of interest (outlined square) was measured. *I*_1_: FI at the dorsal striatum, *I*_2_: FI at the NAc, *I*_3_: FI at the white matter around the midline. Right, comparison of D1R immunoreactivity at the NAc between mice injected with AAV-gRNA (NAc-D1RKO, n = 7 mice) and AAV-Control (Control, n = 5 mice). (C) Heteroduplex cleavage assay. The saCas9-*Drd1a*-gRNA (D1gRNA) showed the capacity of cleaving ON-target (ON) *Drd1a* sequence but not the potential OFF-target (OFF) sites with three mismatches (red letters). Open arrow heads indicate cleavage bands. Control: hrGFP, Neg. Con.: negative control, Pos. Con.: positive control, chr: chromosome. (D) Same as (A) but for data from NAc-D1R-KO (red or orange) and control (black) mice (control: n = 5 mice; NAc-D1R-KO: n = 7 mice). (E) Same as (A) but for data from bReaChES (green or orange) and GFP (black) expressing mice (n = 7 mice for each). Medians of the box plots are shown in magenta or cyan. Open circles and lightly colored lines correspond to individual data. ***p < 0.001, ** p < 0.01, *p < 0.05, paired *t* test (nose length and lateral nose tip movement in (A)), Wilcoxon signed rank test (P(whisk) in (A)) or unpaired *t* test ((B), (D) and (E)).

**Figure S5.**
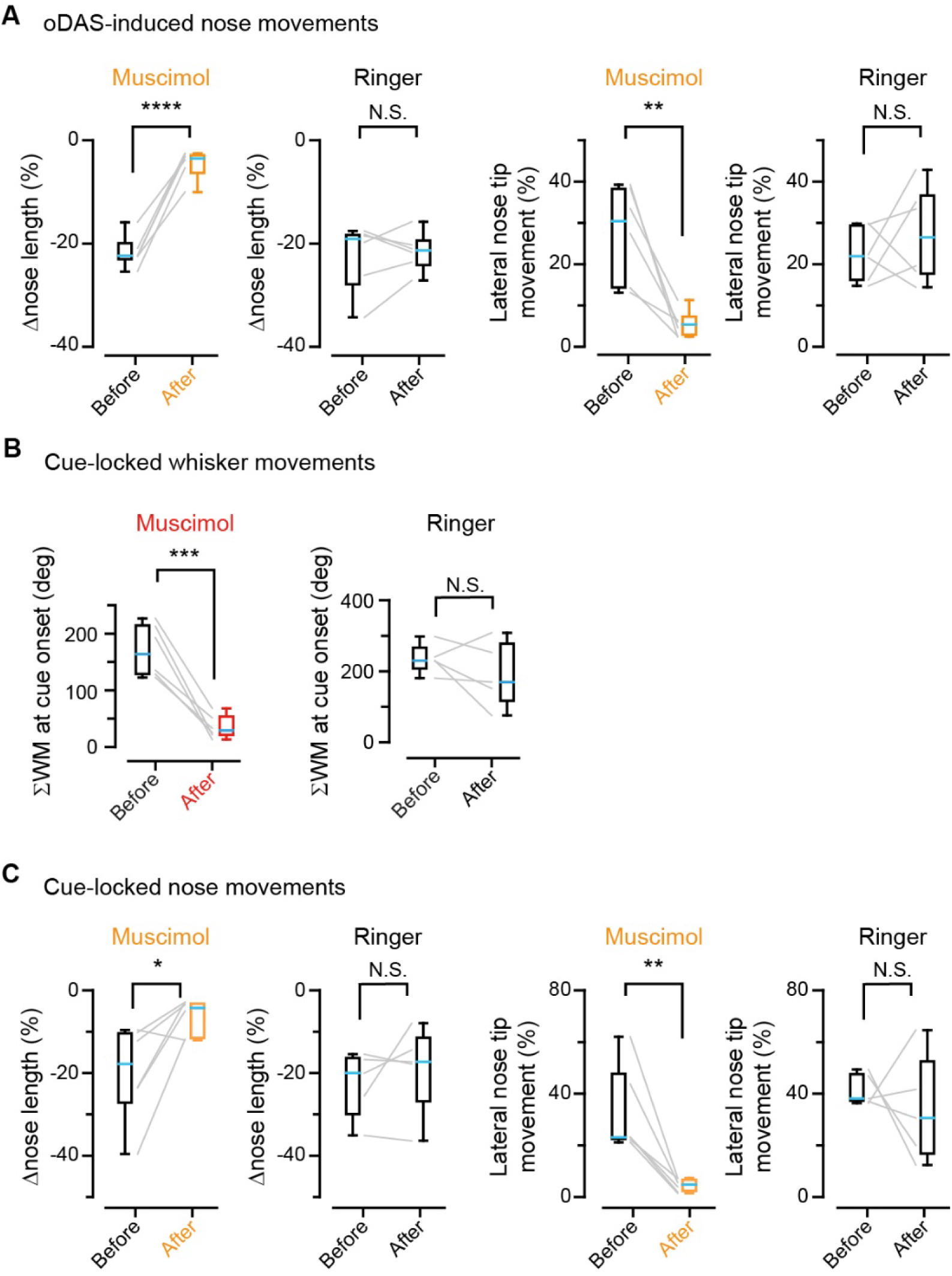
Additional analyses of orofacial movements upon wM1 inhibition. (A) Data for each mouse (thin lines) and box plots for the changes in nose length and the changes in lateral movements of the nose tip during oDAS in trials before and after injection of muscimol or Ringer’s solution (n = 6 mice for each). (B) Same as (A) but for the amount of whisker movements at the cue onset during sound oDAS pairing conditioning on day 3–5 (muscimol, n = 6 mice; Ringer, n = 5 mice). (C) Same as (B) but for nose movement data. Medians of the box plots are shown in cyan. ****p < 0.0001, **p < 0.01, *p < 0.05, N.S., not significant, paired *t* test (Δnose length for Muscimol and lateral nose tip movement for Muscimol and Ringer in (A), all comparisons in (B), and Δnose length for Muscimol and Ringer and lateral nose tip movement for Ringer in (C)) or Wilcoxon signed rank test (Δnose length for Ringer in (A) and lateral nose tip movement for Musimol in (C)).

**Figure S6.**
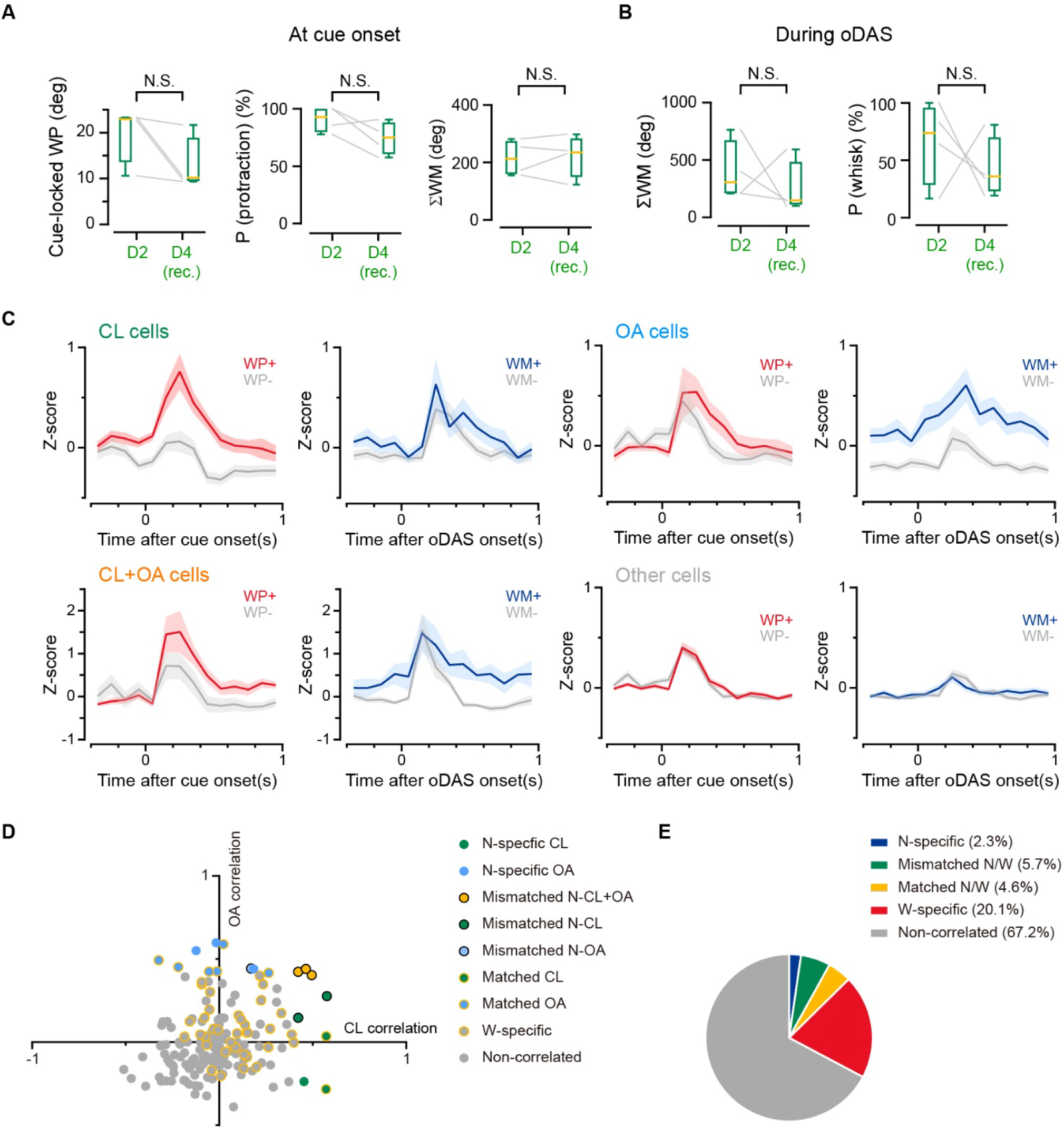
Additional analyses of orofacial movements and their neuronal representation in wM1 in the sound-oDAS pairing experiment. (A) Quantifications of the amplitude (left), probability (middle), and amount (right) of the cue locked whisker protraction (WP). Data on day 2 (D2) and day 4 (D4) of the sound-oDAS pairing conditioning (Figure 7) were obtained from the four DAT-ChR2 mice used for electrophysiological recordings (rec.) at D4. Medians of the box plots are shown in red. N.S., not significant, paired *t* test for P (protraction) and ΣWM or Wilcoxon signed rank test for cue-locked WP. (B) Same as (A) but for the total amount (left) and probability (right) of whisker movements during oDAS. N.S., not significant, paired *t* test for P (whisk) and ΣWM or Wilcoxon signed rank test for ΣWM. (C) Z-scored PSTH of each cellular category (Figure 7E) at the cue onset or upon oDAS. (D) Scatter plot containing 174 neurons from four mice experiencing the sound-oDAS pairing stimulation on the fourth day of conditioning, plotted on the basis of their correlation to the total amount of nose area changes during oDAS (OA correlation) and the amplitude of nose area changes at the cue onset (CL correlation). Subsets of neurons showing significant, positive correlation are colored as indicated in inset. N: nose motion, W: whisker motion. “N-specific” indicates that the neurons’ firings were correlated with nose but not whisker movements. “W-specific” indicates that the neurons’ firings were correlated with whisker but not nose movements. “Matched” means that the neurons exhibited correlation with both nose and whisker movements as the same categories. “Mismatched” means that the neurons showed correlation with both nose and whisker movements but as different categories. (E) Pie chart indicating the proportion of the cell categories.

**Figure S7.**
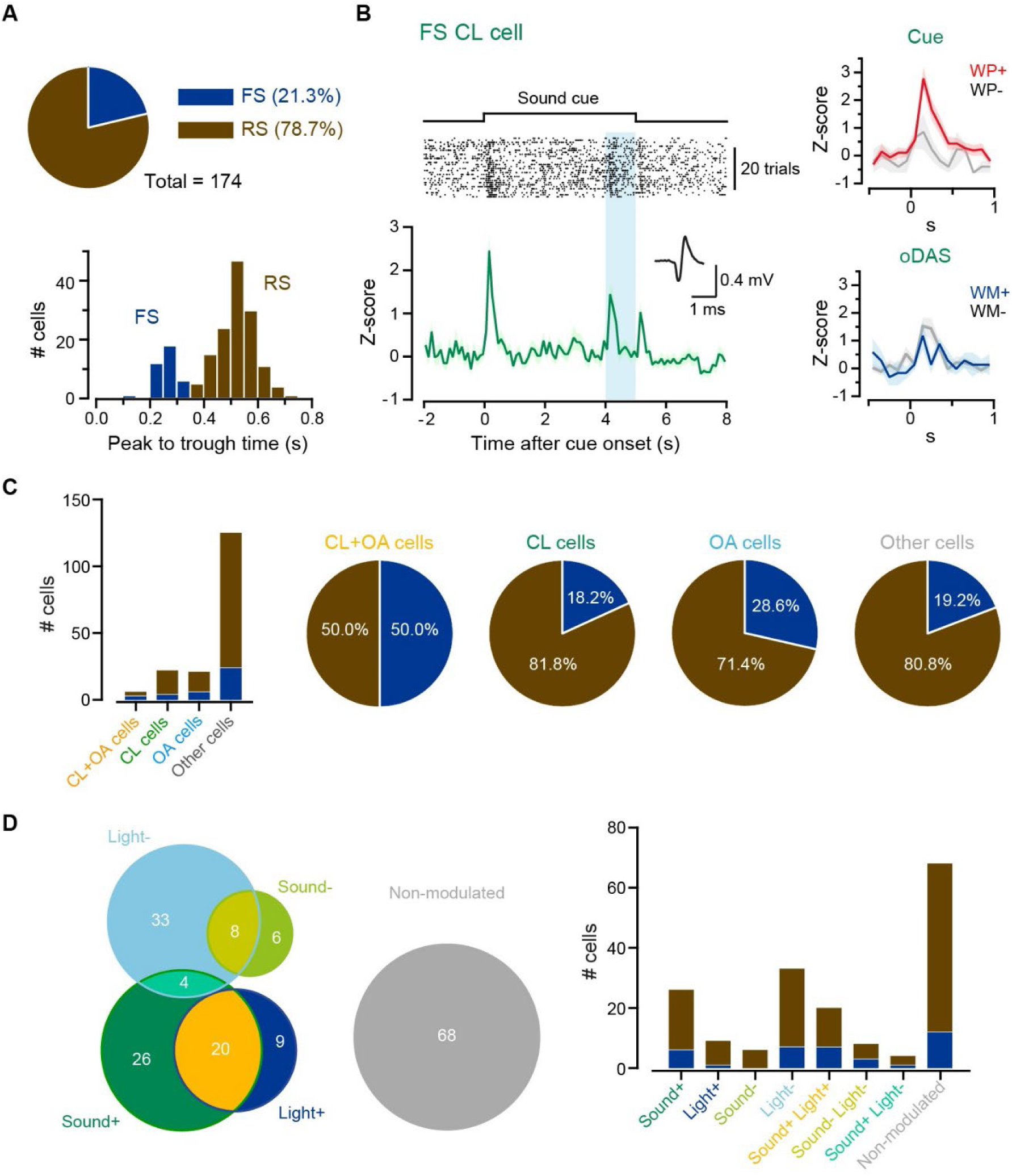
Additional analyses on neuronal activities in wM1 during the sound-oDAS pairing experiment. (A) Top, fraction of fast-spiking (FS, dark blue) and regulate-spiking (RS, copper) cells among recorded units, defined by their peak to trough time of the spike waveform. Bottom, distribution of peak to trough time of recorded units. (B) Left, example raster plot (top) and corresponding Z-scored PSTH (bottom) obtained from a representative FS RE cell. The averaged action potential waveform is shown in the inset. Right, Z-scored PSTH of the same cell at around the cue onset (top) and upon oDAS (bottom), comparing data with or without whisker protraction (WP) at the cue onset or whisker movement (WM) during oDAS. (C) Fraction of FS (dark blue) and RS (copper) neurons among recorded units in the cell categories defined by the correlation with reward-related orofacial movements. (D) Left, among 174 units in wM1 we studied, 103 units (59.2 %) significantly increased (+) or decreased (-) their firing rate at the cue onset (Sound+ or Sound-) and/or during oDAS (Light+ or Light-). Right, fraction of FS (dark blue) and RS (copper) neurons in different cell categories defined by their responsivity (left).

**Figure S8.**
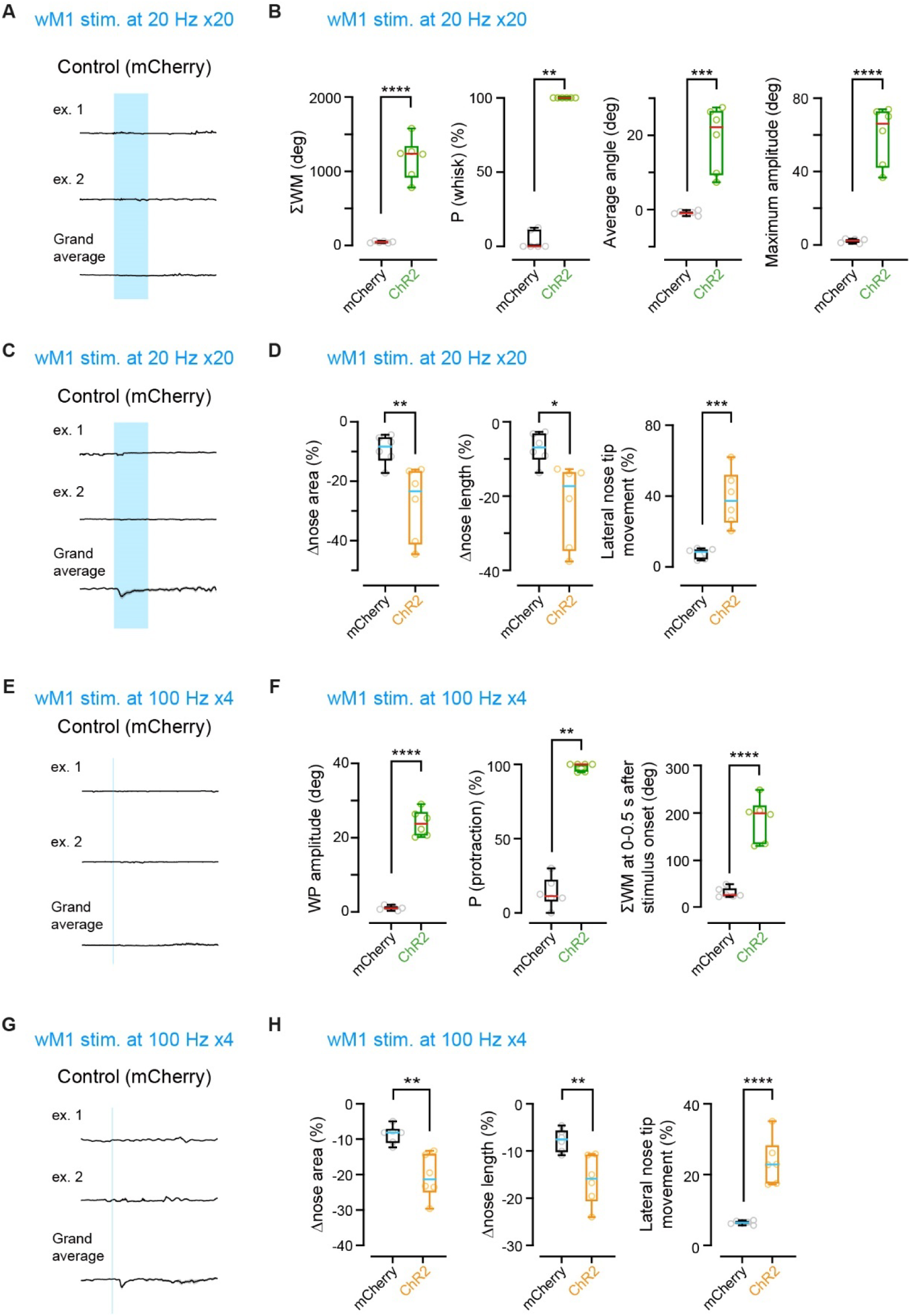
Additional analyses on wM1-driven whisker and nose movements. (A) Example and grand average traces of the whisker position upon photostimulation (5 ms, 20 pulses at 20 Hz) of wM1 expressing mCherry (n = 6 mice). (B) Quantifications for whisker movements evoked by photostimulation of wM1 with 5-ms pulses at 20 Hz for 20 times in mCherry-expressing control mice or ChR2-expressing mice (n = 6 mice for each). (C) Same as (A) but for nose area. (D) Same as (B) but for nose movements. (E–H) Same as (A–D) but with 5-ms pulses at 100 Hz for 4 times. Medians of the box plots are shown in magenta or cyan. ****p < 0.0001, ***p < 0.001, **p < 0.01, *p < 0.05, N.S., not significant, unpaired *t* test, except for Mann-Whitney U test in P(whisk) in (A) and P(protraction) in (C).

## REFERENCES

1. Busse, L., Cardin, J.A., Chiappe, M.E., Halassa, M.M., McGinley, M.J., Yamashita, T., and Saleem, A.B. (2017). Sensation during active behaviors. J. Neurosci. 37, 10826– 10834. 10.1523/JNEUROSCI.1828-17.2017

2. Niell, C.M., and Scanziani, M. (2021). How cortical circuits implement cortical computations: Mouse visual cortex as a model. Ann. Rev. Neurosci. 44, 517–546. 10.1146/annurev-neuro-102320-085825

3. Avitan, L., and Stringer, C. (2022). Not so spontaneous: Multidimensional representations of behaviors and context in sensory areas. Neuron 110, 3064–3075. 10.1016/j.neuron.2022.06.019

4. Stringer, C., Pachitariu, M., Steinmetz, N., Reddy, C.B., Carandini, M., and Harris, K.D. (2019). Spontaneous behaviors drive multidimensional, brainwide activity. Science 364, eaav7893. 10.1126/science.aav7893.

5. Musall, S., Kaufman, M.T., Juavinett, A.L., Gluf, S., and Churchland, A.K. (2019). Single-trial neural dynamics are dominated by richly varied movements. Nat Neurosci 22, 1677–1686. 10.1038/s41593-019-0502-4

6. Steinmetz, N. A., Zatka-Haas, P., Carandini, M., and Harris, K.D. Distributed coding of choice, action and engagement across the mouse brain. Nature 576, 266–273 (2019). 10.1038/s41586-019-1787-x

7. Salkoff, D. B., Zagha, E., McCarthy, E., and McCormick, D.A. (2020). Movement and performance explain widespread cortical activity in a visual detection task. Cereb. Cortex 30, 421–437. 10.1093/cercor/bhz206

8. Tremblay, S., Testard, C., DiTullio, R.W., Inchauspé, J., and Petrides, M. (2023). Neural cognitive signals during spontaneous movements in the macaque. Nat. Neurosci. 26, 295–305. 10.1038/s41593-022-01220-4

9. Bimbard, C., Sit, T.P.H., Lebedeva, A., Reddy, C.B., Harris, K.D., and Carandini, M. (2023). Behavioral origin of sound-evoked activity in mouse visual cortex. Nat. Neurosci. 26, 251–258. 10.1038/s41593-022-01227-x

10. Sachidhanandam, S., Sreenivasan, V., Kyriakatos, A., Kremer, Y., and Petersen, C.C.H. (2013). Membrane potential correlates of sensory perception in mouse barrel cortex. Nat Neurosci 16, 1671–1677. 10.1038/nn.3532

11. Esmaeili, V., Tamura, K., Muscinelli, S.P., Modirshanechi, A., Boscaglia, M., Lee, A.B., Oryshchuk, A., Foustoukos, G., Liu, Y., Crochet, S., et al. (2021). Rapid suppression and sustained activation of distinct cortical regions for a delayed sensory-triggered motor response. Neuron 109, 2183–2201 e2189. 10.1016/j.neuron.2021.05.005.

12. Dominiak, S.E., Nashaat, M.A., Sehara, K., Oraby, H., Larkum, M.E., and Sachdev, R.N.S. (2019). Whisking asymmetry signals motor preparation and the behavioral state of mice. J. Neurosci. 39, 9818–9830. 10.1523/JNEUROSCI.1809-19.2019

13. Coddington, L.T., Lindo, S.E., and Dudman, J.T. (2023). Mesolimbic dopamine adapts the rate of learning from action. Nature 614, 294–302. 10.1038/s41586-022-05614-z

14. Dolensek, N., Gehrlach, D.A., Klein, A.S., and Gogolla, N. (2020). Facial expressions of emotion states and their neuronal correlates in mice. Science 368, 89–94. 10.1126/science.aaz9468

15. Deschênes, M., Moore, J., and Kleinfeld, D. (2012). Sniffing and whisking in rodents. Curr. Opin. Neurobiol. 22, 243–250. 10.1016/j.conb.2011.11.013

16. Welzl, H., and Bureš, J. (1977). Lick-synchronized breathing in rats. Physiol. Behav. 18, 751–753. 10.1016/0031-9384(77)90079-8

17. Moore, J.D., Deschênes, M., Furuta, T., Huber, D., Smear, M.C., Demers, M., and Kleinfeld, D. (2013). Hierarchy of orofacial rhythms revealed through whisking and breathing. Nature 497, 205–210. 10.1038/nature12076.

18. Kurnikova, A., Moore, J.D., Liao, S.M., Deschênes, M., and Kleinfeld, D. (2017). Coordination of Orofacial Motor Actions into Exploratory Behavior by Rat. Curr. Biol. 27, 688–696. 10.1016/j.cub.2017.01.013

19. Schultz, W., Dayan, P., and Montague, P.R. (1997). A neural substrate of prediction and reward. Science 275, 1593–1599. 10.1126/science.275.5306.1593.

20. Berridge, K.C., and Robinson, T.E. (1998). What is the role of dopamine in reward: hedonic impact, reward learning, or incentive salience? Brain Res Brain Res Rev 28, 309–369. 10.1016/s0165-0173(98)00019-8

21. Berridge, K.C., and Kringelbach, M.L. (2015). Pleasure systems in the brain. Neuron 86, 646–664. 10.1016/j.neuron.2015.02.018

22. Tsai, H.C., Zhang, F., Adamantidis, A., Stuber, G.D., Bonci, A., de Lecea, L., and Deisseroth, K. (2009). Phasic firing in dopaminergic neurons is sufficient for behavioral conditioning. Science 324, 1080–1084. 10.1126/science.1168878

23. Adamantidis, A.R., Tsai, H.C., Boutrel, B., Zhang, F., Stuber, G.D., Budygin, E.A., Tourino, C., Bonci, A., Deisseroth, K., and de Lecea, L. (2011). Optogenetic interrogation of dopaminergic modulation of the multiple phases of reward-seeking behavior. J. Neurosci. 31, 10829–10835. 10.1523/JNEUROSCI.2246-11.2011

24. Pascoli, V., Hiver, A., Van Zessen, R., Loureiro, M., Achargui, R., Harada, M., Flakowski, J., and Lüscher, C. (2018). Stochastic synaptic plasticity underlying compulsion in a model of addiction. Nature 564, 366–371. 10.1038/s41586-018-0789-4

25. Burgess, C.P., Lak, A., Steinmetz, N.A., Zatka-Haas, P., Bai Reddy, C., Jacobs, E.A.K., Linden, J.F., Paton, J.J., Ranson, A., Schroder, S., et al. (2017). High-yield methods for accurate two-alternative visual psychophysics in head-fixed mice. Cell Rep. 20, 2513–2524. 10.1016/j.celrep.2017.08.047

26. Cohen, J.Y., Haesler, S., Vong, L., Lowell, B.B., and Uchida, N. (2012). Neuron-type specific signals for reward and punishment in the ventral tegmental area. Nature 482, 85–88. 10.1038/nature10754

27. Tian, J., and Uchida, N. (2015). Habenula lesions reveal that multiple mechanisms underlie dopamine prediction errors. Neuron 87, 1304–1316. 10.1016/j.neuron.2015.08.028

28. Starkweather, C. K., Babayan, B. M., Uchida, N., and Gershman, S.J. (2017). Dopamine reward prediction errors reflect hidden-state inference across time. Nat. Neurosci. 20, 581–589 10.1038/nn.4520

29. Brecht, M., Schneider, M., Sakmann, B., and Margrie, T.W. (2004). Whisker movements evoked by stimulation of single pyramidal cells in rat motor cortex. Nature 427, 704–710. 10.1038/nature02266

30. Matyas, F., Sreenivasan, V., Marbach, F., Wacongne, C., Barsy, B., Mateo, C., Aronoff, R., and Petersen, C.C.H. (2010). Motor control by sensory cortex. Science 330, 1240–1243. 10.1126/science.1195797

31. Sreenivasan, V., Karmakar, K., Rijli, F.M., and Petersen, C.C.H. (2015). Parallel pathways from motor and somatosensory cortex for controlling whisker movements in mice. Eur. J. Neurosci. 41, 354–367. 10.1111/ejn.12800

32. Sreenivasan, V., Esmaeili, V., Kiritani, T., Galan, K., Crochet, S., and Petersen C.C.H. (2016). Movement initiation signals in mouse whisker motor cortex. Neuron 92, 1368–1382. 10.1016/j.neuron.2016.12.001

33. Mercer Lindsay, N., Knutsen, P.M., Lozada, A.F., Gibbs, D., Karten, H.J., and Kleinfeld, D. (2019). Orofacial movements involve parallel corticobulbar projections from motor cortex to trigeminal premotor nuclei. Neuron 104, 765–780. 10.1016/j.neuron.2019.08.032

34. Hill, D.N., Curtis, J.C., Moore, J.D., and Kleinfeld, D. (2011). Primary motor cortex reports efferent control of vibrissa motion on multiple timescales. Neuron 72, 344–356. 10.1016/j.neuron.2011.09.020

35. Friedman, W.A., Zeigler, H.P., and Keller, A. (2012). Vibrissae motor cortex unit activity during whisking. J. Neurophysiol. 107, 551–563. 10.1152/jn.01132.2010

36. Gerdjikov, T.V., Haiss, F., Rodriguez-Sierra, O.E., and Schwarz, C. (2013). Rhythmic whisking area (RW) in rat primary motor cortex: an internal monitor of movement related signals? J. Neurosci. 33, 14193–14204. 10.1523/JNEUROSCI.0337-13.2013

37. Yamashita, T., and Petersen, C.C.H. (2016). Target-specific membrane potential dynamics of neocortical projection neurons during goal-directed behavior. eLife 5, e15798. 10.7554/eLife.15798

38. Mathis, A., Mamidanna, P., Cury, K.M., Abe, T., Murthy, V.N., Mathis, M.W., and Bethge, M. (2018). DeepLabCut: markerless pose estimation of user-defined body parts with deep learning. Nat. Neurosci. 21, 1281–1289. 10.1038/s41593-018-0209-y

39. Dodson, P. D., Dreyer, J.K., Jennings, K.A., Syed, E.C.J., Wade-Martins, R., Cragg, S.J., Bolam, J.P., and Magill P.J. (2016). Representation of spontaneous movement by dopaminergic neurons is cell-type selective and disrupted in parkinsonism. Proc. Natl. Acad. Sci. USA 113, E2180–E2188. 10.1073/pnas.1515941113

40. Howe, M.W., and Dombeck, D.A. (2016). Rapid signalling in distinct dopaminergic axons during locomotion and reward. Nature 535, 505–510. 10.1038/nature18942

41. Gunaydin, L.A., Grosenick, L., Finkelstein, J.C., Kauvar, I.V., Fenno, L.E., Adhikari, A., Lammel, S., Mirzabekov, J.J., Airan, R.D., Zalocusky, K.A., et al. (2014). Natural neural projection dynamics underlying social behavior. Cell 157, 1535–1551. 10.1016/j.cell.2014.05.017

42. Cui, W., Aida, T., Ito, H., Kobayashi, K., Wada, Y., Kato, S., Nakano, T., Zhu, M., Isa, K., Kobayashi, K., et al. (2020). Dopaminergic signaling in the nucleus accumbens modulates stress-coping strategies during inescapable stress. J. Neurosci. 40, 7241– 7254. 10.1523/JNEUROSCI.0444-20.2020

43. Bindra, D., and Campbell, J.F. (1967). Motivational effects of rewarding intracranial stimulation. Nature 215, 375–376 10.1038/215375a0

44. Clarke, S., and Trowill, J.A. (1971). Sniffing and motivated behavior in the rat. Physiol. Behav. 6, 49–52. 10.1016/0031-9384(71)90013-8

45. Kepecs, A., Uchida, N., and Mainen, Z.F. (2006). The sniff as a unit of olfactory processing. Chem Senses 31, 167–179. 10.1093/chemse/bjj016

46. Saunders, B.T., Richard, J.M., Margolis, E.B., and Janak, P.H. (2018). Dopamine neurons create Pavlovian conditioned stimuli with circuit-defined motivational properties. Nat. Neurosci. 21, 1072–1083. 10.1038/s41593-018-0191-4

47. Welker, W.I. (1964). Analysis of sniffing of the albino rat. Behaviour 22, 223–244. 10.1163/156853964X00030

48. Bayer, H. M. & Glimcher, P. W. Midbrain dopamine neurons encode a quantitative reward prediction error signal. Neuron 47, 129–141 (2005).

49. Petersen, C.C.H. (2014). Cortical control of whisker movement. Annu Rev Neurosci 37, 183–203. 10.1146/annurev-neuro-062012-170344

50. McElvain, L.E., Friedman, B., Karten, H.J., Svoboda, K., Wang, F., Deschênes, M., and Kleinfeld, D. (2018). Circuits in the rodent brainstem that control whisking in concert with other orofacial motor actions. Neuroscience 368, 152–170. 10.1016/j.neuroscience.2017.08.034

51. Takatoh, J., Prevosto, V., Thompson, P.M., Lu, J., Chung, L., Harrahill, A., Li, S., Zhao, S., He, Z., Golomb, D., et al. The whisking oscillator circuit. Nature 609, 560–568 (2022). 10.1038/s41586-022-05144-8

52. Patriarchi, T., Cho, J.R., Merten, K., Howe, M.W., Marley, A., Xiong, W-H., Folk, R.W., Broussard, G.J., Liang, R., Jang, M.J., et al. (2018). Ultrafast neuronal imaging of dopamine dynamics with designed genetically encoded sensors. Science 360, eaat4422. 10.1126/science.aat4422

53. Pachitariu, M., Steinmetz, N.A., Kadir, S.N., Carandini, M., and Harris, K.D. (2016). Fast and accurate spike sorting of high-channel count probes with KiloSort. 30th Conference on Neural Information Processing. In Advances in Neural Information Processing 29 (NIPS 2016), D. Lee, M. Sugiyama, U. Luxburg, I. Guyon, R. Garnett, eds. (NeurIPS Proceedings)

54. Madisen, L., Mao, T., Koch, H., Zhuo, J.M., Berenyi, A., Fujisawa, S., Hsu, Y.W., Garcia 3rd, A.J., Gu, X., Zanella, S., et al. (2012). A toolbox of Cre-dependent optogenetic transgenic mice for light-induced activation and silencing. Nat. Neurosci. 15, 793–802 10.1038/nn.3078

55. Matsubara, T., Yanagida, T., Kawaguchi, N., Nakano, T., Yoshimoto, J., Sezaki, M., Takizawa, H., Tsunoda, S.P., Horigane, S.-I., Ueda, S., et al. (2021). Remote control of neural function by X-ray-induced scintillation. Nat. Commun. 12, 4478. 10.1038/s41467-021-24717-1

